# The GR2D2 Estimator for the Precision Matrices

**DOI:** 10.1101/2022.03.22.485374

**Authors:** Dailin Gan, Guosheng Yin, Yan Dora Zhang

## Abstract

Biological networks are important for the analysis of human diseases, which summarize the regulatory interactions and other relationships between different molecules. Understanding and constructing networks for molecules, such as DNA, RNA and proteins, can help elucidate the mechanisms of complex biological systems. The Gaussian Graphical Models (GGMs) are popular tools for the estimation of biological networks. Nonetheless, reconstructing GGMs from high-dimensional datasets is still challenging. Current methods cannot handle the sparsity and high-dimensionality issues arising from datasets very well. Here we developed a new GGM, called the GR2D2 (Graphical *R*^2^-induced Dirichlet Decomposition) model, based on the R2D2 priors for linear models. Besides, we provided a data-augmented block Gibbs sampler algorithm. The R code is available at https://github.com/RavenGan/GR2D2. The GR2D2 estimator shows superior performance in estimating the precision matrices compared to existing techniques in various simulation settings. When the true precision matrix is sparse and of high dimension, the GR2D2 provides the estimates with smallest information divergence from the underlying truth. We also compare the GR2D2 estimator to the graphical horseshoe estimator in five cancer RNA-seq gene expression datasets grouped by three cancer types. Our results show that GR2D2 successfully identifies common cancer pathways and cancer-specific pathways for each dataset.

## 1 Introduction

Biological networks, which capture the regulatory interactions or other relationships between molecules, such as DNA, RNA and protiens, are important for us to understand the mechanisms of complicated biological systems [3, 13]. The genetic regulatory networks, for instance, have played important roles of analysing gene expression data, for they can capture the complex gene-gene interactions and make the inferences of the dependency structures of genes possible [39, 40]. The Gaussian Graphical Models (GGMs) have been a popular tool in genomics to describe the functional dependencies between genes of interests in graphs, since these models can circumvent indirect associations between genes by evaluating conditional dependencies in multivariate Gaussian distributions [17, 18, 30, 39]. GGMs have also been successfully applied to different scenarios to infer the genetic regulatory networks [7, 19, 24, 29, 31, 41]. In GGMs, we assume that the multivariate vector of genes follows a multivariate normal distribution with a particular structure of the inverse of the covariance matrix, called the precision matrix. Under this assumption, the conditional independence of two genes is equivalent to a zero value of the corresponding element in the precision matrix. The graph generated by GGMs can provide lucid interpretations of the relationship between genes in which nodes represent genes and edges between them represent interactions. The set of edges, which decides the network topology, can also be used to generate hypotheses about genetic interactions [18].

However, reconstructing GGMs from high-dimensional datasets remain a difficult task. The standard estimation of the partial correlation of genes either estimates the precision matrix or the parameters in *p* least squares regression problems, where *p* is the number of genes. If the sample size *n* is much smaller than the number of genes *p*, both methods are inappropriate [16]. Possible alternatives are either using penalization or the Bayesian framework to estimate the precision matrix. Consider *n* samples from a *p*-dimensional multivariate normal distribution with zero mean and a *p* × *p* covariance matrix Ω^−1^, which is

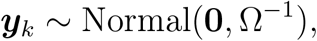

where *k* = 1, 2, …, *n* and Ω is the precision matrix. The *i, j*-th off-diagonal element of Ω is the negative of the partial covariance between features *i* and *j*, and the *i*-th feature is regressed on all the other features [26]. Besides, under the multivariate normal assumption, zero off-diagonal elements in Ω are conditionally independent given the remaining features.

One major challenge in estimating precision matrices is that the number of parameters increases quadratically with the number of features, leading to a sparse matrix. One approach to deal with sparsity is to penalized the likelihood functions. In graphical lasso [9], they maximize the penalized likelihood

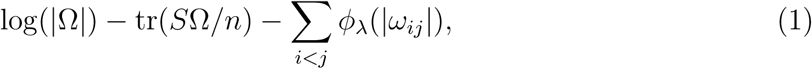

where 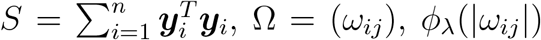 is the *l*_1_ penalty, and *λ* is a tuning parameter. A Bayesian version of the graphical lasso [36] is proposed afterwards, and the goal is to maximize a posterior estimate of Ω under the prior

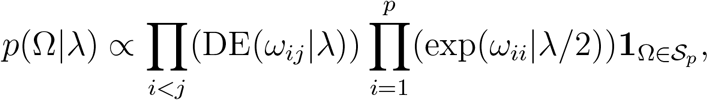

where DE(*x* | *λ*) denotes the double exponential distribution with rate *λ*, exp(*x* | *λ*) denotes the exponential distribution with rate *λ*, and *𝒮*_*p*_ is the space of *p* × *p* positive definite matrices.

The smoothly clipped absolute deviation (SCAD) penalty [8] was also introduced to estimate the precision matrix, in which they maximized the penalized likelihood in equation 1 where the penalty term has the first-order derivative

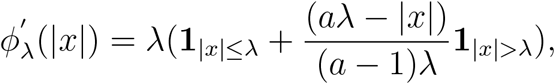

with *a* > 2 and *λ* > 0. However, the performance of this method relies on the choice of tuning parameters, which cannot be easily implemented in reality.

The graphical horseshoe estimator employs a Bayesian framework and puts horseshoe priors on the off-diagonal elements of precision matrices, and puts uninformative prior on the diagonal elements [20]. The model is summarized in the following

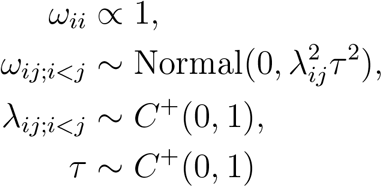

where *C*^+^(0, 1) denotes a half-cauchy random variable with density *p*(*x*) ∝ (1 + *x*^2^)^−1^; *x* > 0. The distinctive scale parameter *λ*_*ij*_ is treated as a local shrinkage parameter, and the scale parameter τ is shared by all dimensions, treated as a global shrinkage parameter.

The graphical lasso, Bayesian graphical lasso, and the graphical SCAD face the problem of accumulated estimation errors due to increasing number of parameters [20], while the graphical horseshoe estimator shows weak performance in high-dimensional settings when the number of features *p* is much larger than the sample size *n*. Here, we propose an alternative Bayesian estimator called the GR2D2 (Graphical *R*^2^-induced Dirichlet Decomposition) estimator that can handle sparse and high-dimensional issues faced by other methods. The R2D2 shrinkage priors have shown promising performance in high-dimensional linear regression models, having greater concentration near the origin and heavier tails than current shrinkage priors [42]. We extended the R2D2 priors from linear models to graphical models and our simulation results show that the GR2D2 estimators achieved the state-of-art performance in sparse and high-dimensional settings, compared with the graphical lasso and the graphical horseshoe estimator. We further compare the GR2D2 estimator to the graphical horseshoe estimator in five cancer RNA-seq gene expression datasets grouped by three cancer types. Our results show that the graphical horseshoe estimator always gives highly densely connected networks and it is very hard to identify relevant gene sets for a specific cancer pathway. GR2D2 outputs sparse networks with clear structures and it successfully identifies common cancer pathways and cancer-specific pathways.

The remaining sections are organized as follows. We outlines the GR2D2 estimator with a full Gibbs sampler for efficient sampling. Next we present our simulation studies to show the efficiency of the GR2D2 followed by real data analysis.

## 2 Methods

For Gaussian graphical models, it is usually assumed that the patterns of variation in expression of a gene can be predicted by those of a small subset of other genes, which leads to a sparse structure in the precision matrix [40]. For such precision matrix, we need a shrinkage method that is able to give zero or very small estimate for the zero elements. Besides, this method should also be able to sustain the nonzero values and shrink them as little as possible. We propose the GR2D2 estimator to achieve this goal.

### 2.1 The GR2D2 Hierarchical Model

The GR2D2 model puts R2D2 priors on the off-diagonal elements of the precision matrix. The element-wise priors are specified for *i, j* = 1, …, *p* as follows

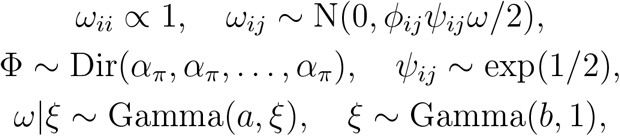

where N(0, 1) denotes a normal random variable with mean 0 and variance 1; Dir(*α*_*π*_, *α*_*π*_, …, *α*_*π*_) denotes a vector of Dirichlet distributed random variables with a parameterized vector ***α*** = [*α*_*π*_, *α*_*π*_, …, *α*_*π*_]^*T*^ ; and the vectors ***α*** and Φ are of length *p*^2^. However, in the following notations, we use Φ as a symmetric matrix of dimension *p* × *p* with entries *ϕ*_*ij*_ and use ***ϕ*** as a row or column vector of Φ. Further, let exp(1/2) denote an exponential random variable with rate 1/2; let Gamma(1, 1) denote a Gamma random variable with shape 1 and rate 1. The distinctive scale parameters *ϕ*_*ij*_ and *ψ*_*ij*_ on each dimension are referred to as the local shrinkage parameters, and the scale parameter *ω* shared by all dimensions is referred to as the global shrinkage parameter. Note that *ξ* is used as an auxiliary parameter to help sample *ω*. Since conditional independence of each entry is guaranteed in precision matrices given the remaining entries under the multivariate normal assumption, the prior on Ω under the GR2D2 model can be written as the product of off-diagonal entries

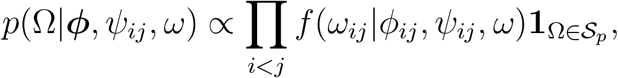

where *f*(*ω*_*ij*_ *ϕ*_*ij*_, *ψ*_*ij*_, *ω*) stands for the normal density function of *ω*_*ij*_ given parameters *ϕ*_*ij*_, *ψ*_*ij*_, *ω*, and 𝒮_*p*_ is the space of *p*× *p* positive definite matrices.

In high-dimensional precision matrix estimation using GR2D2, the global shrinkage parameter *ω* adapts to the sparsity of the entire matrix Ω and shrinks the estimates of the off-diagonal elements toward zero. On the flip side, the local shrinkage parameters, *ϕ*_*ij*_ and *ψ*_*ij*_ for *i* < *j*, preserve the magnitude of nonzero off-diagonal elements, and ensure that the element-wise biases are not very large.

### 2.2 A Data-Augmented Block Gibbs Sampler

Posterior samples under the GR2D2 hierarchical model are drawn by an augmented block Gibbs sampler, utilizing the scheme proposed by [22] for linear regression. In each iteration, each column and row of Ω and Λ = (*ϕ*_*ij*_*ψ*_*ij*_) are partitioned from a *p*× *p* matrix of parameters and updated in a block. Then the global shrinkage parameter *ω* and its auxiliary variable *ξ* are updated.

Given data *Y*_*n*×*p*_ and shrinkage parameters, we provide necessary derivation of the posterior distribution of the precision matrix as follows. Define the matrices

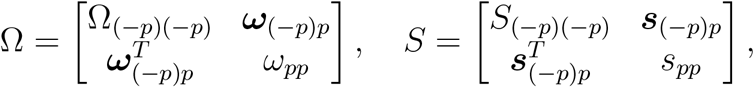

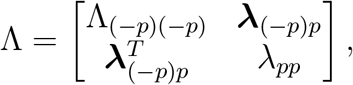

where the subindex _(−*p*)_ denotes the set of all indices except for p, and the matrix Λ has entries *λ*_*ij*_ = *ϕ*_*ij*_*ψ*_*ij*_. Diagonal elements of Λ_(−*p*)(−*p*)_ are set to be 1. The posterior distribution of Ω is

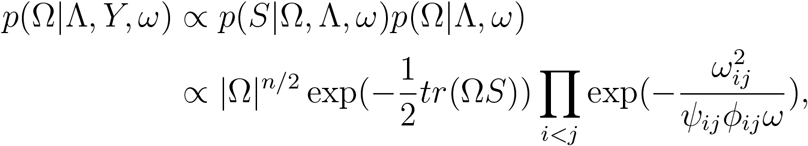

where *S* = *Y* ^*T*^ *Y* ∼ *W*_*p*_(Ω^−1^, *n*) and *W* stands for a Wishart distribution. After some calculation, the full conditional of the last column of Ω is

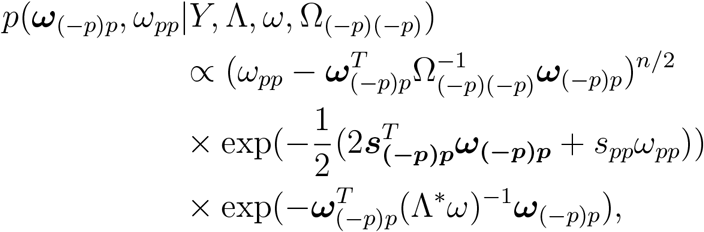

where 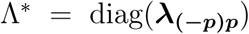. Let 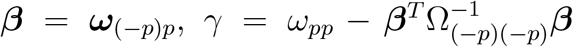, and 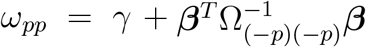. The Jacobian of this transformation is a constant, and the full conditional of *β* and *γ* is

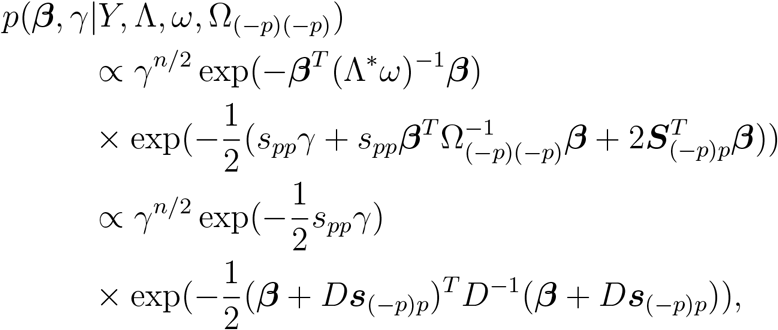

where *D* = (2(Λ^∗^*ω*)^−1^ + *s*_*pp*_Ω_(−*p*)(−*p*)_)^−1^. The above formula can be summarized as

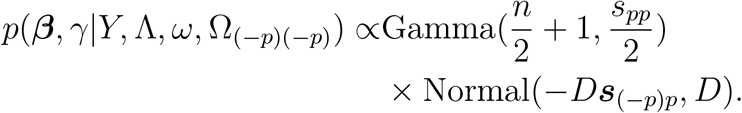

Hence, the posterior distribution of the last row and column of Ω is obtained. Since the derivation of the posterior distributions of *ψ*_*ij*_, *ω, ξ* and ***ϕ*** is standard, we summarize them in the sampling procedure section with a proposed algorithm structure.

### 2.3 Gibbs sampling procedures with initial settings

Based on the results derived in the previous section, we provide the initial settings used in our sampling procedures. We use the same initial settings as [20] and [42], and set *p* to be the number of rows (or columns) in *S* = *Y* ^*T*^ *Y*, Ω = *I*_*p*×*p*_, Σ = *I*_*p*×*p*_, Φ = **1**_*p*×*p*_, Ψ = **1**_*p*×*p*_, *ω* = 1, ξ = 1, *b* = 0.5, *r* = 1, *C* = 1, 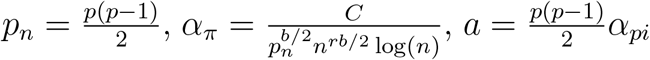, Λ = Φ ⊙ Ψ, where Φ has entries of *ϕ*_*ij*_, Ψ has entries of *ψ*_*ij*_. The Gibbs sampling procedures are as follows:

a. Set initial values as above.
b. Choose the *p*-th column and do the following until the matrix Ω is updated.
  (b.1) Sample *γ*|*Y*, Λ, *ω*, Ω_(−*p*)(−*p*)_ ∼ Gamma(shape = 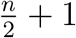, rate = 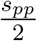).
  (b.2) Sample ***β***|*Y*, Λ, *ω*, Ω_(−*p*)(−*p*)_ ∼ Normal(−*D****s***_(−*i*)*i*_, *D*), where *D* = (2(Λ^∗^*ω*)^−1^ + s_*pp*_Ω_(−*p*)(−*p*)_)^−1^, Λ^∗^ = diag(***λ***_(−*p*)*p*_), and *λ*_*ij*_ = *ϕ*_*ip*_*ψ*_*ip*_.
  (b.2) Sample 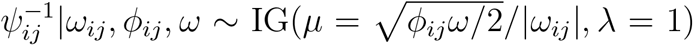, and then take the reciprocal to get ψ_*ij*_. IG stands for an inverse Gamma distribution.
c. Sample *ω*|Ω, Λ, *ξ* ∼ giG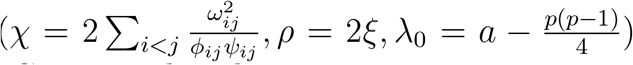 where giG stands for a generalized inverse Gaussian distribution.
d. Sample *ξ*|*ω* ∼ Gamma(shape = *a* + *b*, rate = *ω* + 1)
e. Sample ***ϕ***|Ω, *ψ, *ω*, ξ*. Based on our derivation, if 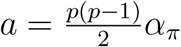 one can draw 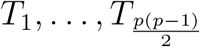 independently with 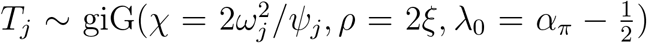. Then set *ϕ*_*j*_ = *T*_*j*_/*T* with 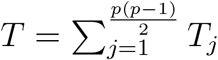
f. Repeat steps (b)-(g) until convergence.

The algorithm 1 of the GR2D2, along with simulation examples, is available at this github website.

## 3 Simulation Studies

In the graphical horseshoe estimator [20], they compared the performance of the graphical horseshoe with the graphical SCAD[8], frequentist graphical lasso [9] and the Bayesian graphical lasso [36] using diverse assessment criteria and different structures of the precision matrices. Their results showed that the graphical horseshoe estimator achieved the best performance. Therefore, we only compare the GR2D2 estimator with the graphical horseshoe estimator and the graphical lasso, where the graphical lasso works as the baseline.

### 3.1 Simulation Setting and Assessment Criterion

We consider six precision matrix structures with different pairs of features and observations. These pairs are (*p, n*) = (100, 50), (150, 50), (200, 50), and (250, 50). The reason of this setting is that we intend to compare the GR2D2 and other estimators when the ratio *p*/*n* increases, which corresponds to high-dimensional settings. The diagonal entries of the precision matrix *ω*_0_ is 1 for all six structures, while the off-diagonal entries depend on one of the following structures. We take the pair (*p, n*) = (100, 50) as an example.

#### Algorithm 1 The GR2D2 Sampler

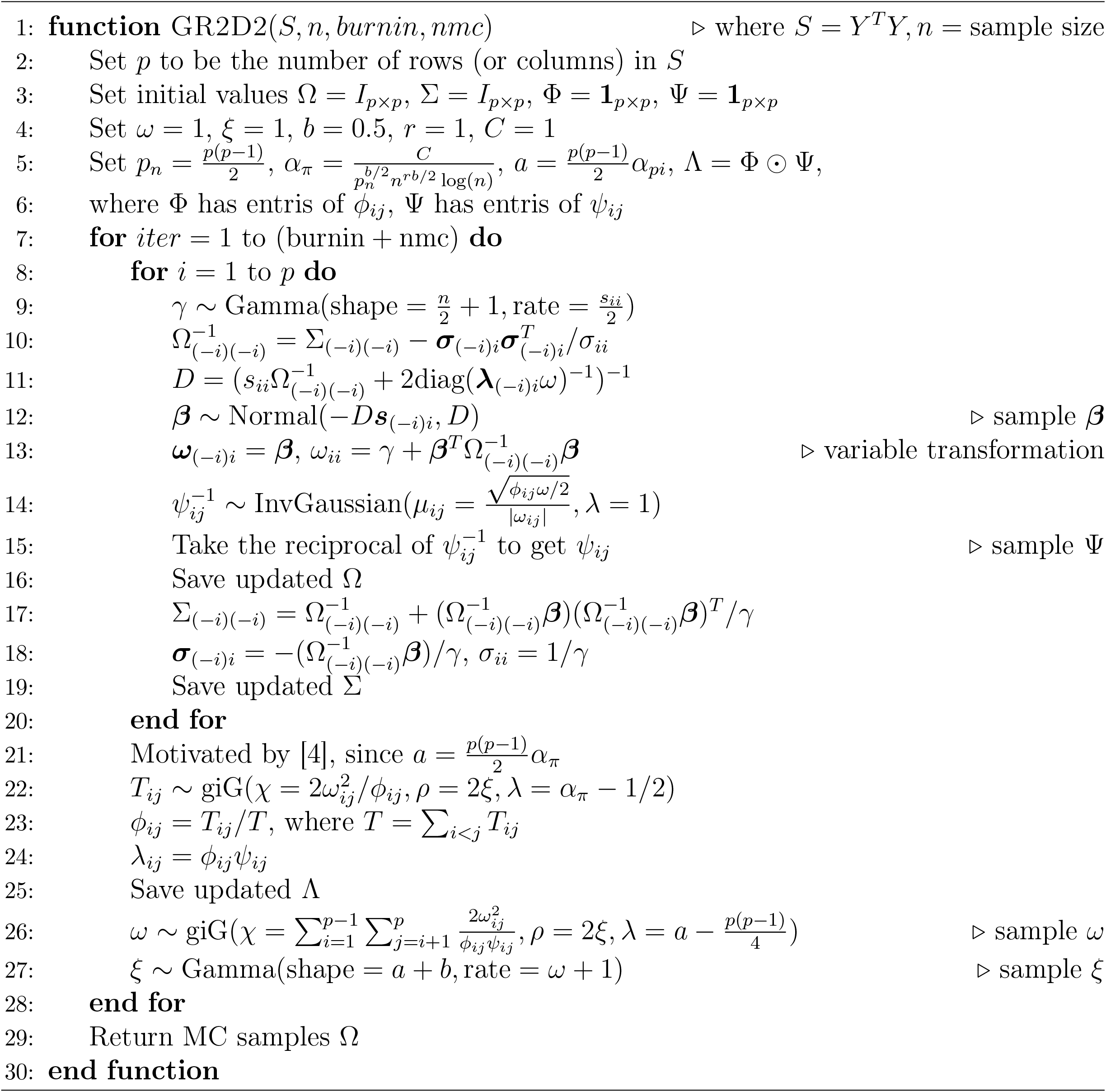

#### Hubs

The rows or columns are partitioned into disjoint groups 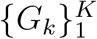 Each group has a row k where off-diagonal elements are taken to be *ω*_*ik*_ = 0.25 corresponding to negative partial correlations for *i* ∈ *G*_*k*_ and *ω*_*ij*_ = 0 otherwise, and we allow four members within each group. We consider 10 groups and 10 members within each group, giving 40 nonzero off-diagonal elements in Ω_0_. The topological structure of this pattern is like a star with four members connected with one member at the center.

#### Cliques positive

The rows or columns are partitioned into disjoint groups and {*ω*_*ij*_ : *i, j* ∈ *G*_*k*_, *i* ≠ *j*} are set to -0.45 corresponding to positive partial correlation. We also use 10 groups in this setting but only also three members within each group resulting in 30 nonzero off-diagonal elements in Ω_0_. The topological structure of this pattern is like a small circle with only three points connected with each other.

#### Cliques negative

The rows or columns are partitioned into disjoint groups and {*ω*_*ij*_ : *i, j* ∈ *G*_*k*_, *i* ≠ *j*} are set to 0.75 corresponding to positive partial correlation. Other settings are the same as those in cliques positive.

#### Scale-free positive

We use the function barabasi.game from the igraph R package to generate a scale-free graph according to the Barabasi-Albert model (BA model) [2]. 100 (n = 100) vertices are generated with the power of the preferential attachment equal to 1 (power = 1). We set m = 1 to add one edge in each time step, resulting in a sparse network structure. The arguments n, power and m are used in the function barabasi.game. We then turn the graph into an adjacency matrix, which is treated as the precision matrix in our estimation. The off-diagonal elements are set to be -0.15, corresponding to the positive partial correlations.

#### Scale-free negative

The settings in this structure are identical to those in scale-free positive except that the off-diagonal elements are set to be 0.15, corresponding to the negative partial correlations.

#### Scale-free half

The settings in this structure are identical to those in scale-free positive except that we randomly sample half of the off-diagonal elements to be 0.15 and the other half to be -0.15.

We choose the hubs, cliques positive and negative structures since they are commonly used structures to verify the performance of graphical models [10]. One more structure, random, mentioned in [10], is not used here since the generated samples from the random structure may not always be positive definite and hence not invertible. We use three scale-free graph structures based on the BA model. This model generates graphs whose vertex connectivities follow a scale-free power-law distribution. This power-law distribution has been found in many natural and man-made networks [6]. Hence, we apply this scale-free structure to generate graphs that may represent realistic network structures.

The results of four pairs of observations and features and six structures are summarized in Figures 1 and 2, and in Supplementary Information Tables S1-S8. We also include the results of three scale-free structures using (*p, n*) = (250, 30) in Table S9 to show the performance of graphical models when the sample size is small. Considering the variation in the network topology [11], in Tables S10 and S11, we generated 30 different networks for Hubs, Cliques positive and Cliques negative structures with 30 different non-zero *ω*’s and (*p, n*) = (100, 50) and (150, 50). (For Hubs, *ω* is ranging from 0.11 to 0.4 with step size 0.01; for Cliques positive, *ω* is ranging from -0.40 to -0.1 with step size 0.01; for Cliques negative, *ω* is ranging from 0.11 to 0.4 with step size 0.01) In these figures, we report the results of Frobenius norm, Stein’s loss, precision, and recall, while in tables we also report false positive rate (FPR), accuracy, and F scores with corresponding standard deviation (s.d.).

**Figure 1:**
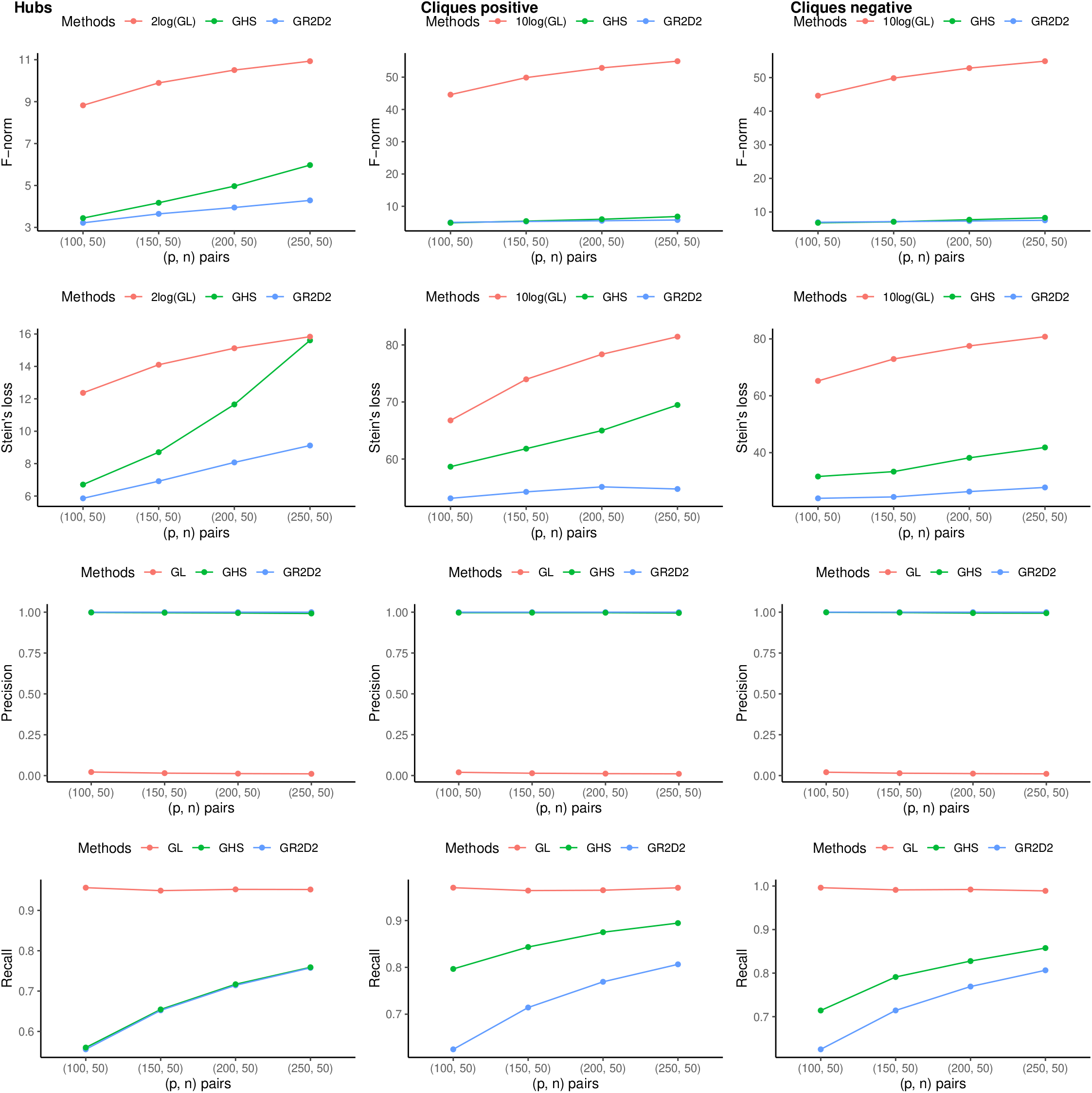
Frobenius norm (F-norm), Mean Stein’s loss, precision, and recall of precision matrix estimates for hubs, cliques positive and negative network structures over 30 datasets generated by multivariate normal distribution with the sample size *n* = 50 and number of features *p* = (100, 150, 200, 250). The precision matrix is estimated by frequentist graphical lasso (GL), the graphical horseshoe (GHS), and the GR2D2. We use 2log(GL), 10log(GL) and 10log(GL) for the GL estimator in hubs, cliques positive and negative respectively when comparing F-norm and Stein’s loss. For F-norm, GR2D2 performs better than or comparably well to GHS while GL performs poorly. For Stein’s loss, GR2D2 consistently shows the best performance. For precision scores, GR2D2 and GHS show comparable good performance while GL provides the worst precision scores. For recall scores, GR2D2 and GHS show comparable performance in hubs structure while GHS performs better in cliques structure. GL performs the best regarding to the recall scores. Overall, GR2D2 shows the best estimation results according to F-norm and Stein’s loss and comparable results to GHS in variable selection under hubs, cliques positive and negative network structures

**Figure 2:**
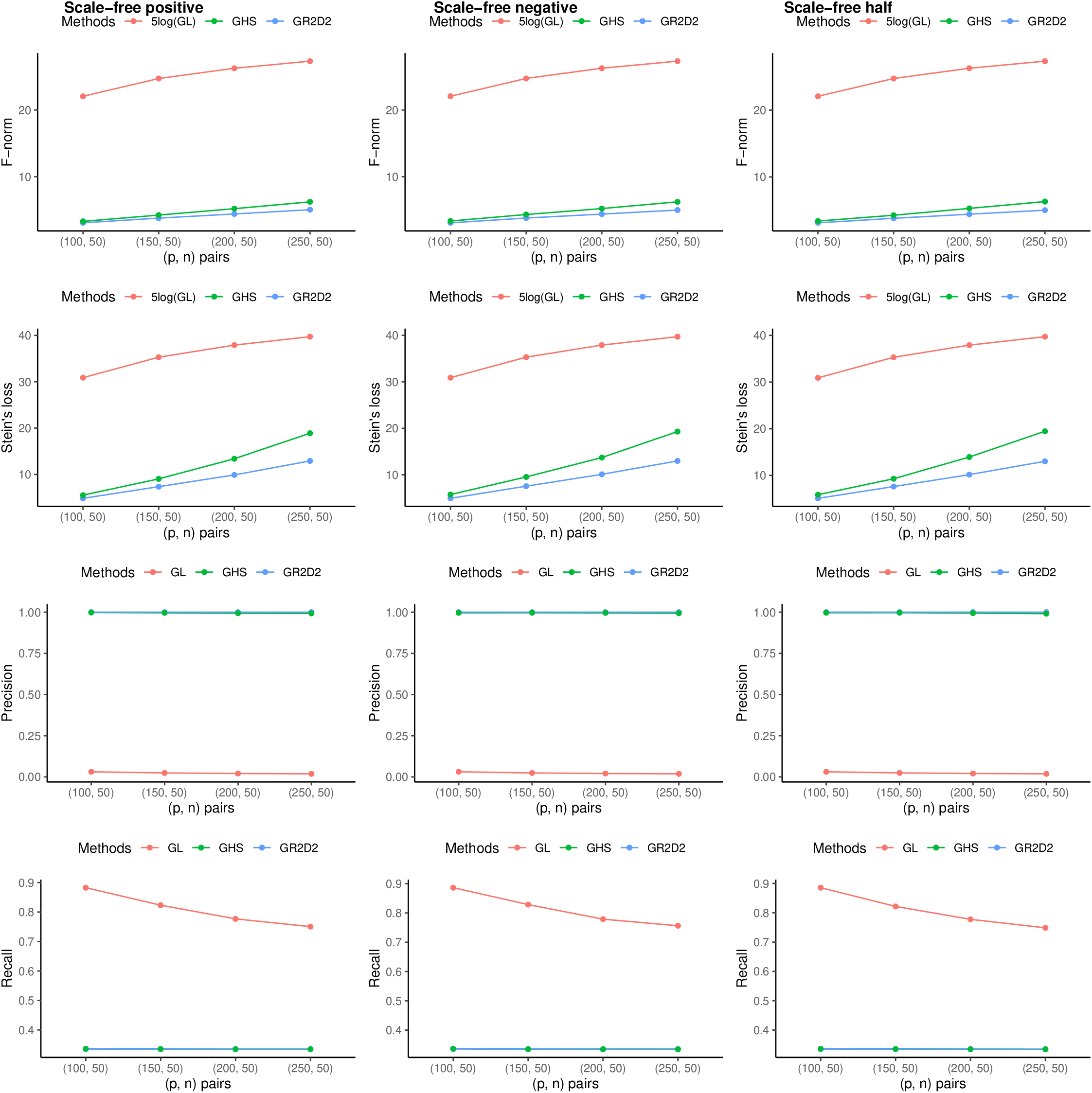
Frobenius norm (F-norm), Mean Stein’s loss, precision, and recall of precision matrix estimates for scale-free positive, negative and half network structures over 30 datasets generated by multivariate normal distribution with the sample size *n* = 50 and number of features *p* = (100, 150, 200, 250). The precision matrix is estimated by frequentist graphical lasso (GL), the graphical horseshoe (GHS), and the GR2D2. We use 5log(GL) for the GL estimator in scale-free positive, negative and half when comparing F-norm and Stein’s loss. For F-norm and Stein’s loss, GR2D2 consistently shows the best performance. For precision scores, GR2D2 and GHS show comparable good performance while GL provides the worst precision scores. For recall scores, GR2D2 and GHS show comparable performance while GL performs the best. Overall, GR2D2 shows the best estimation results according to F-norm and Stein’s loss and comparable results to GHS in variable selection under scale-free positive, negative and half network structures.

For each choice of Ω_0_, 30 datasets are generated and Ω is estimated using the graphical lasso, the graphical horseshoe and GR2D2 estimators. The graphical horseshoe estimator is implemented in MATLAB (2022a), while the graphical lasso is implemented using the package “glasso” in Rstudio (version 4.1.2). For the graphical horseshoe and the GR2D2 estimators, we drop the initial 500 burnin samples and keep the following 500 MCMC samples with convergence. We also use the median of the remaining 500 samples as our estimate for the graphical horseshoe and GR2D2.

The Stein’s loss of the estimated precision matrix 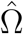 is equal to 2 times the Kullback-Leibler divergence of 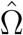 from Ω_0_, with the formula

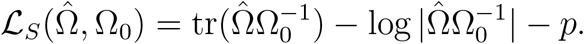

The Frobenius norm between 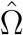 and Ω_0_ is

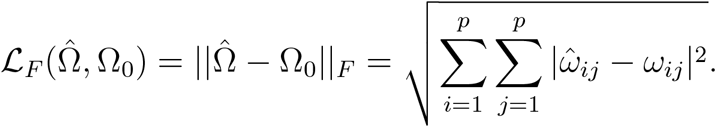

Since the graphical horseshoe and the GR2D2 are shrinkage methods that do not estimate elements to be exactly equal to zero, we use the 95% quantile intervals in the MCMC samples for variable selection. This is to say, if the 95% quantile intervals of an off-diagonal entry of Ω does not contain zero, this entry is considered as a discovery and the median values across the MCMC samples are used as the estimate. If the discoveries identified in 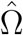 have the same signs as those in Ω_0_, we call them true positive discoveries and use them to calculate the precision, recall, positive rate, accuracy and F scores. For each measurement, we report the mean and s.d. computed over 30 datasets.

### 3.2 Estimation

Based on Figures 1 and 2, the GR2D2 estimates always have the smallest Steins’ loss and most of the time have the smallest Frobenius norm (F norm). When the ratio *p*/*n* is 2, the graphical horseshoe estimators show better performance than GR2D2 in F norm in the structures cliques positive and cliques negative; and when the ratio between *p* and *n* is 3, the graphical horseshoe estimators show better performance than GR2D2 in F norm in the pattern cliques negative only. When *p*/*n* is 4 and 5, the GR2D2 shows consistent superior performance to the graphical horseshoe in all six structures. The GR2D2 is expected to perform well when the precision matrix is sparse and the absolute values of scaled nonzero elements are large as mentioned in the work of [43]. Besides, the GR2D2 is able to deal with high dimensional datasets and always provides the best results as shown in *p*/*n* = 4 and *p*/*n* = 5. These results show that in both commonly used network structures and scale-free network structures, GR2D2 performs overall the best.

Besides, it could be noted that as the ratio between p and n increases, the Stein’s loss and F norm increase accordingly, which is expected since when the datasets become sparser and the dimension becomes higher, it is harder to estimate the precision matrices. Considering the standard deviation across 30 datasets, the GR2D2 always gives 0 values, indicating that the estimation of GR2D2 could provide the most concentrated estimation results with least variation, as shown in Tables S1-S11. For the graphical horseshoe estimator, it shows relatively small variation while the graphical lasso provides the biggest variations.

### 3.3 Variable Selection

As shown in Figures 1 and 2, for the precision, GR2D2 always gives 1, indicating the all the positive observations are true observations. The graphical horseshoe provides results close to 1, showing comparably good results, while the graphical lasso provides quite small precision scores. For the scores of recall, the graphical lasso always gives the best recall scores while the results of graphical horseshoe and the GR2D2 improve as the ratio between *p* and *n* increases. Besides, the graphical horseshoe estimator most of the time gives better scores than the GR2D2. For FPR, as shown in Tables S1-S11, the GR2D2 always gives 0 values, indicating that there are 0 false positive result across all 30 datasets. The graphical horseshoe estimators provide results quite close to zero across 30 datasets, showing comparably good results to the GR2D2. However, the graphical lasso always provides the worst FPR scores, indicating that this method always provides false positive estimations. For the accuracy scores, both the graphical horseshoe and the GR2D2 show comparably good results with values more than 0.99, while the graphical lasso provides poorer accuracy results. One point that is worth mentioning is that since the precision matrices are quite sparse, there are always much less observations than zero values, resulting in unbalanced datasets. Therefore, the accuracy scores may not be a good measurement to assess the performance of three estimation methods. Next we turn to F scores, which can cooperate the results of precision and recall, giving measurement free from the effects of unbalanced datasets. According to results of F scores, the graphical horseshoe method and GR2D2 show comparable results in the six structures, while the graphical lasso performs poorly. We also put the formulae of precision, recall, FPR, accuracy and F scores in the Supplementary Information Section 2. Overall, both GR2D2 and the graphical horseshoe show promising abilities to correctly identify the true observations in variable selection under our simulation settings, while the graphical lasso shows weak performance.

## 4 Real data analysis

### 4.1 Datasets and pre-processing steps

Five RNA-seq gene expression datasets of cancerous tissues are obtained from the TCGA database via TCGAbiolinks (v. 2.25.2) [37]. These datasets are breast invasive carcinoma (BRCA) with 1094 cancer patients, kidney renal clear cell carcinoma (KIRC) with 533 cancer patients, kidney renal papillary cell carcinoma (KIRP) with 290 cancer patients, lung adenocarcinoma (LUAD) with 515 cancer patients, and lung squamous cell carcinoma (LUSC) with 502 cancer patients. We use the Kyoto Encyclopedia of Genes and Genomes database to download three cancer pathways [15], which are breast cancer (hsa05224), renal cell carcinoma (hsa05211), and non-small-cell lung cancer (hsa05223). In the following analysis, the cancer gene expression datasets from a specific organ are pooled to form one dataset for that organ, resulting in three datasets, BRCA, KIRC_KIRP, and LUAD_LUSC.

For the pre-processing steps, since the downloaded datasets have been normalized by fragments per kilobase of transcript per million mapped reads, we replace 0 by 1 and add 1 to all the non-zero values before the log-transformation. Genes specific to each pathway are extracted and pooled to form the subset of genes for all three datasets. Finally, we have three datasets with 212 genes each before applying graphical models.

We assume the gene expressions of individuals are identically distributed with a multi-variate normal distribution. Simular to the steps we use for simulation studies, we drop the initial 500 burnin samples and keep the following 500 MCMC samples. We use 99% quantile intervals in the MCMC samples for variable selection to get sparse networks. If 0 is not included in the quantile interval, we treated it as a discovery. Based on the results in our simulation studies, since both GR2D2 and the graphical horseshoe show comparably good performance in variable selection, we compare both methods in our real data analysis.

## Results

The inferred graphs by GR2D2 of datasets BRCA, KIRC_KIRP and LUAD_LUSC are shown in Figures 3, 4, 5. We also provide the inferred graph by the graphical horseshoe of the LUAD_LUSC dataset in Supplementary Information Figures S1. With 99% quantile intervals, the graphical horseshoe method identifies 1704 edges for the LUAD_LUSC dataset, resulting in a densely connected graph. This densely connected graph indicate that the graphical horseshoe estimator fails to select the most relevant edges for the dataset, and it becomes hard for the users to identify useful information from the network structure. Similar results on datasets BRCA and KIRC_KIRP can be expected. On the other hand, the GR2D2 method identifies 94 edges for the BRCA dataset, 76 edges for the KIRC_KIRP dataset, and 108 edges for the LUAD_LUSC dataset, resulting in sparse graphs for all three datasets.

**Figure 3:**
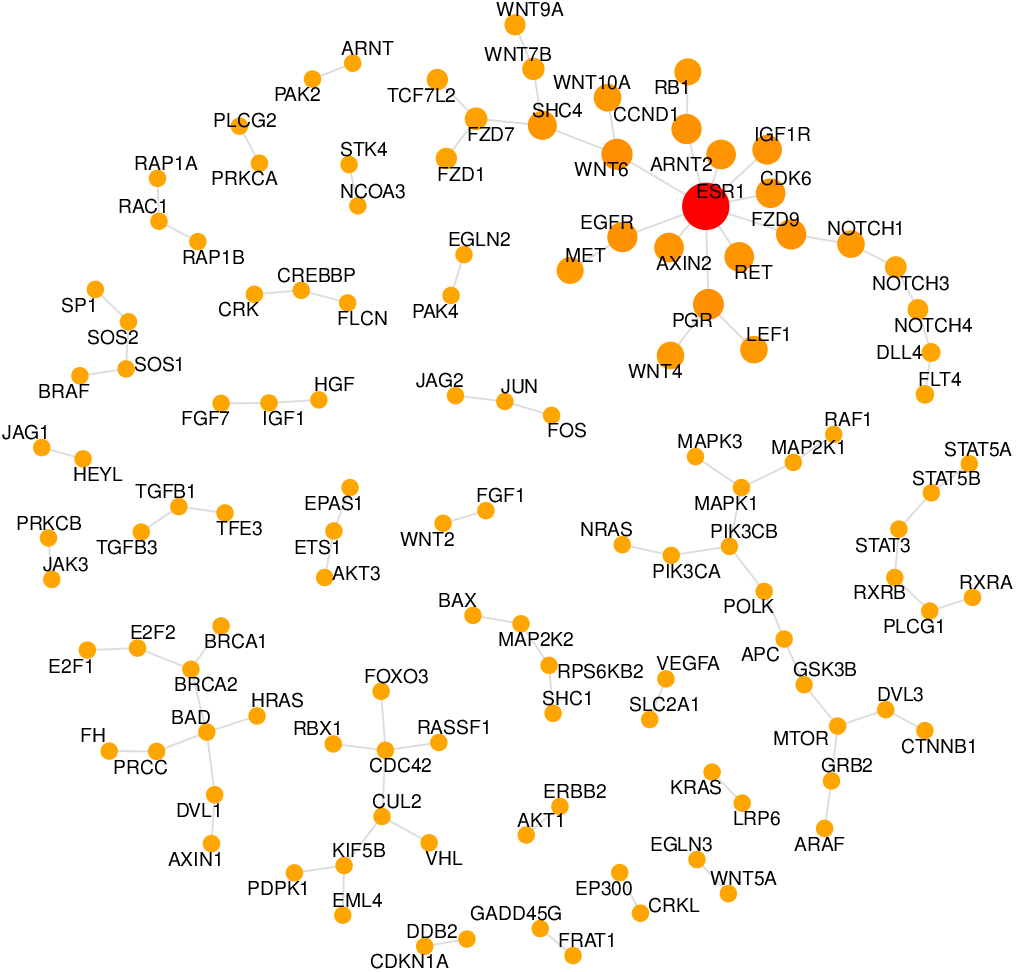
The inferred graph for the BRCA dataset by the GR2D2 estimator. The size of node is proportional to the degrees of edges. At the center to the right part and the top right part of this figure, four pathways can be identified, which are MAPK, Wnt, Notch and ESR1 signaling pathways. MAPK and Wnt pathways are commonly found in many cancers while Notch and ESR1 have special relevance to the breast cancer.

**Figure 4:**
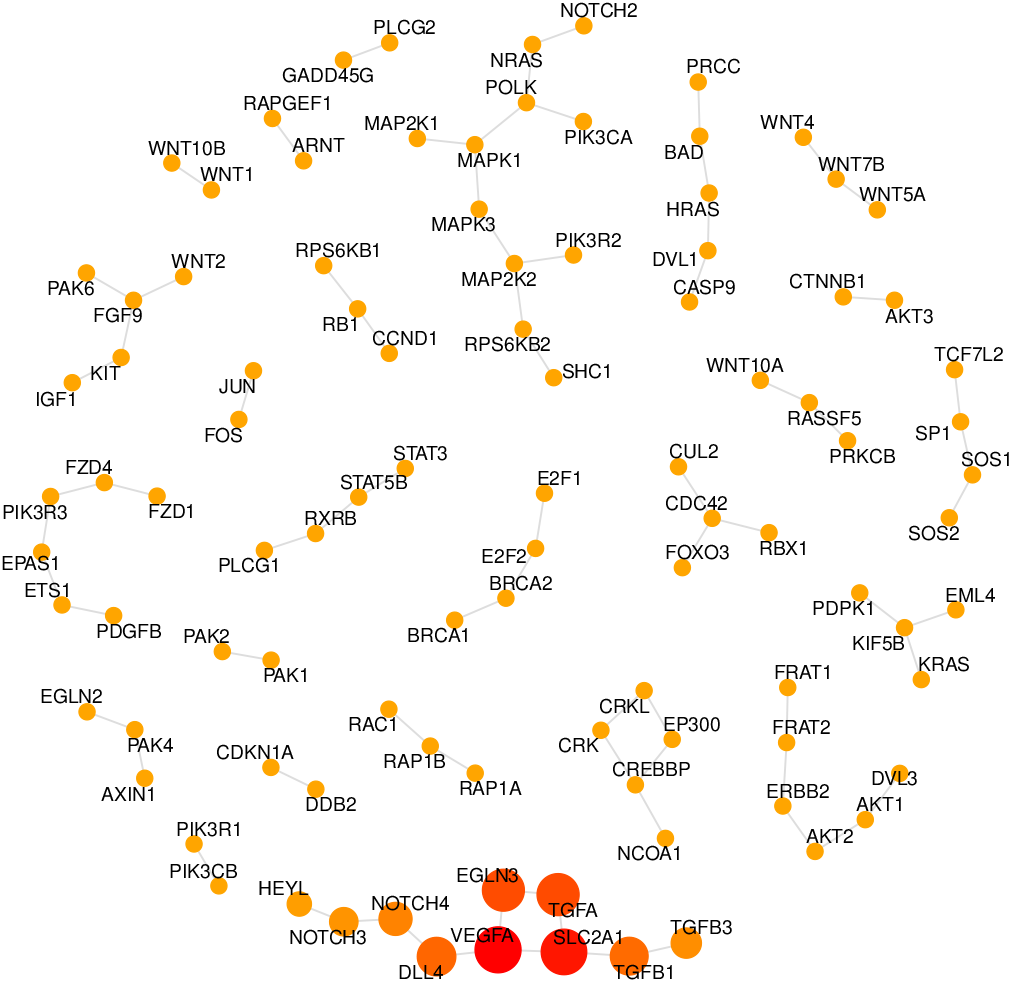
The inferred graph for the KIRC_KIRP dataset by the GR2D2 estimator. The size of node is proportional to the degrees of edges. At the upper part and the lower part of the figure, genes from MAPK and HIF-1 signaling pathways can be found. HIF-1 pathway has special relevance to kidney disease while the MAPK pathway is related to many human cancers.

**Figure 5:**
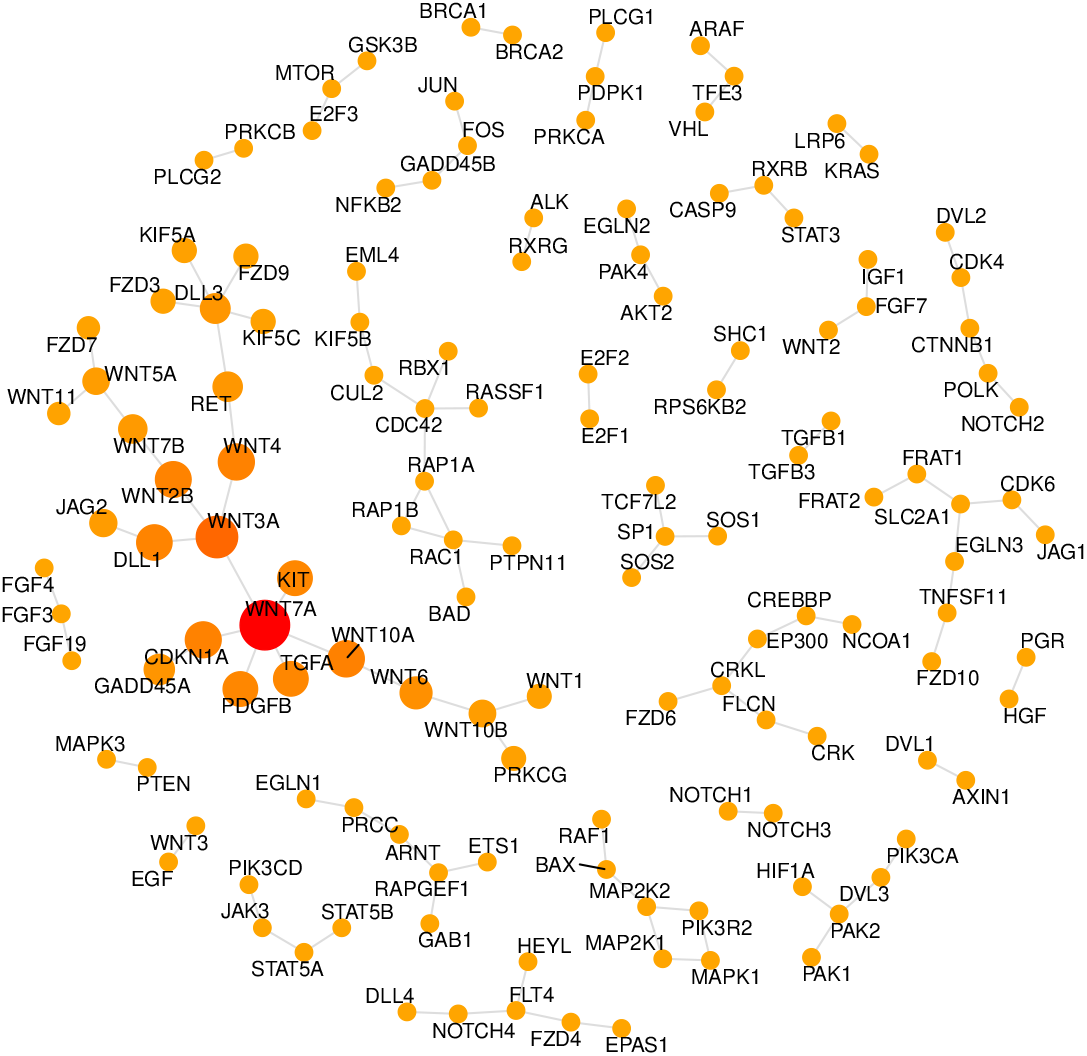
The inferred graph for the LUAD_LUSC dataset by the GR2D2 estimator. The size of node is proportional to the degrees of edges. At the center to the left part, lower left part and lower right part of the figure, Wnt, JAK-STAT and MAPK signaling pathways can be identified. MAPK and Wnt pathways are commonly found in many cancers while JAK-STAT has been found to be related to the progression of non-small cell lung cancer. The KIT gene at the left part of the figure is also identified, which shows special relevance to lung cancer.

We further scrutinize the graphs inferred by GR2D2 and check whether important cancer pathways have been identified. At the center to the right of Figure 3, the upper part of Figure 4, and the bottom of Figure 5, we notice a community of genes formed by RAF1, MAPK1, MAP2K1, MAPK2K2, and MAPK3, which correspond to the genes in the MAPK signaling pathway. This pathway comprises several necessary signaling components that play a role in tumorigenesis, and the alternation of it has been reported in human cancer as a result of abnormal activation of receptor tyrosine kinases [28, 35]. Besides, this pathway is considered as potential therapeutic targets for cancer treatment and several inhibitors targeting this pathway have been developed [28]. At the top right part of Figure 3 and the left part of Figure 5, there are two communities of genes, most of which come from the Wnt signaling pathway. This pathway regulates many cellular processes, like cell proliferation, differentiation and cell fate, and aberrant Wnt signaling pathway is a hallmark of many cancers, including breast cancer [25, 38] and non-small cell lung cancer [33].

We then check pathways and genes that seem only appear in specific cancer types. At the top right of Figure 3, besides genes from the Wnt siganling pathway, genes (NOTCH1/3/4, DLL4, FLT4) from the Notch signaling pathway and genes (ESR1, CCND1) from the ESR1 signaling pathway are identified. Activated Notch signaling pathway and upregulation of tumor-promoting Notch target genes have been reported in human breast cancer and this pathway has been implicated in both the development and progression of breast cancer [1, 27, 34]. The ESR1 ESR1 signaling pathway has been found to be responsible for the luminal substype of breast cancer and the mutation of the gene ESR1 has been reported in approximately 70% of breast cancer patients [14]. The ESR1 gene is also treated as a biomarker in breast cancer [5]. At the bottom of Figure 4, genes (VEGFA, TGFA, SLC2A1, TGCB1, and TGFB3) from the HIF-1 signaling pathway are identified. The activation of this pathway has special relevance to the kidney disease [12, 23] and we cannot find it on the other two graphs. At the left bottom of Figure 5, genes (JAK3, STAT5A and STAT5B) from the JAK-STAT signaling pathway are identified. This pathway has been found to be related to the progression of non-small cell lung cancer [44]. We further notice that the gene KIT at the center to the left part of Figure 5 is also identified. The mutation of KIT has special relevance to the lung cancer [21, 32]. Based on above findings, we can conclude that the GR2D2 estimator identifies important cancer pathways and also cancer-specific pathways for each cancer type.

## 5 Discussion

In high-dimensional data analysis, how to estimate the precision matrix is always an attractive challenge under multivariate Gaussian assumption. In this work, we proposed the GR2D2 estimator with easy implementation by a full Gibbs sampler. By using a prior with high density near the origin, a combination of an exponential distributed parameter and a Dirichlet distributed parameter on each dimension, the GR2D2 hierarchical model provides estimates close to the true distribution in Kullback-Leibler divergence and with the smallest bias for nonzero elements among current published methods. Simulation studies also confirm that the GR2D2 outperforms alternative methods like the graphical horseshoe and the graphical lasso in different settings. Furthermore, real data analysis shows that the GR2D2 estimator can capture important cancer-specific genetic interactions and have the ability to provide insights for future experiment design. Therefore, we may conclude that the GR2D2 could provide the best precision matrix estimation in sparse and high-dimensional settings, and infer meaningful genetic relationships.

## 6 Data availability

Data used in this study were downloaded from public databases that have been described in the main text.

## 7 Supplementary information and code

Supplementary information are available online with detailed derivation of GR2D2. The source code is available at https://github.com/RavenGan/GR2D2.

## Supplementary Information

### 1. A Data-Augmented Block Gibbs Sampler

Posterior samples under the graphical R2D2 hierarchical model are drawn by an augmented block Gibbs sampler, utilizing the scheme proposed by [3] for linear regression. In each iteration, each column and row of Ω and Λ = (*ϕ*_*ij*_*ψ*_*ij*_) are partitioned from a *p* × *p* matrix of parameters and updated in a block. Then the global shrinkage parameter *ω* and its auxiliary variable *ξ* are updated.

The following part derives the posterior distribution of the precision matrix. Given data *Y*_*n*×*p*_ and the shrinkage parameters, we provide the step-by-step derivation in the following. Define the following matrices

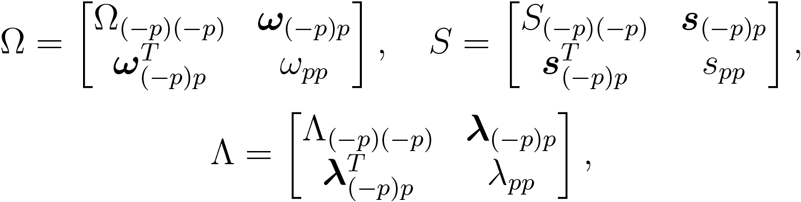

where (−*p*) denotes the set of all indices except for p, and Λ_(−*p*)(−*p*)_ and ***λ***_(−*p*)*p*_ have entries *ϕ*_*ij*_*ψ*_*ij*_. Diagonal elements of Λ_(−*p*)(−*p*)_ are set to be 1. The posterior distribution of Ω is

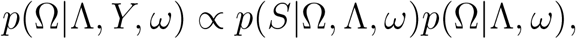

where *S* = *Y* ^*T*^ *Y* ∼ *W*_*p*_(Ω^−1^, *n*) has the Wishart density function

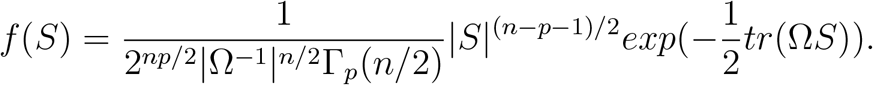

Then, the posterior distribution of Ω can be written as

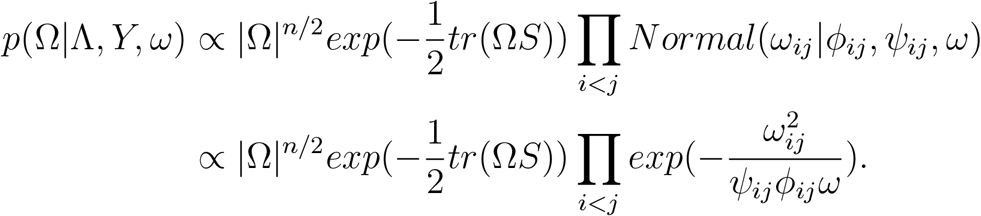

By block determinant, we have

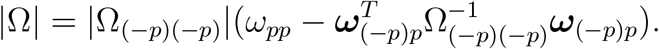

Therefore, we have

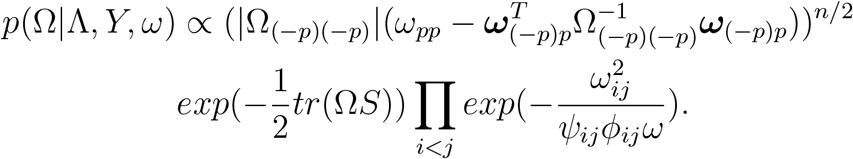

Then we consider *tr*(*S*Ω),

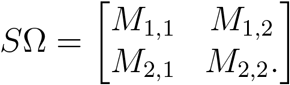

where

- 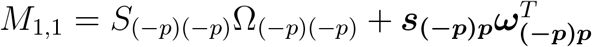
- *M*_1,2_ = *S*_(−*p*)(−*p*)_***ω***_**(*−p*)*p***_ + *ω*_*pp*_***s***_**(*−p*)*p***_
- 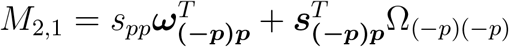
- 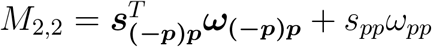

Note that

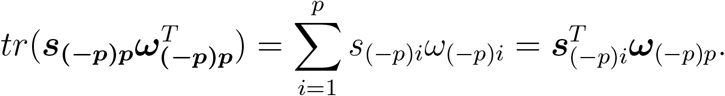

Hence, we obtain

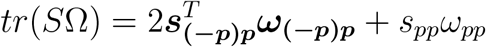

If we only consider the last column

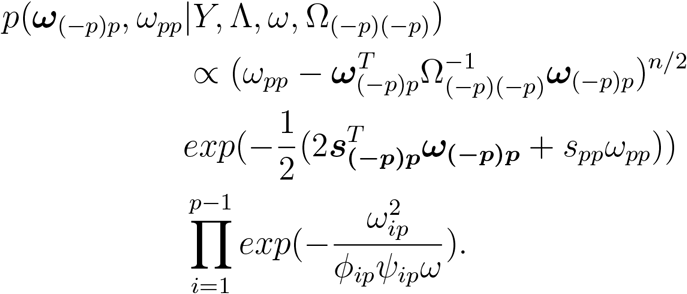

Note that

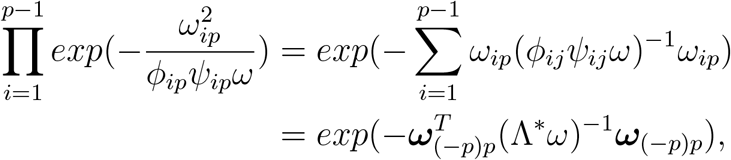

where Λ^∗^ = *diag*(***λ***_**(*−p*)*p***_).

Therefore, we have

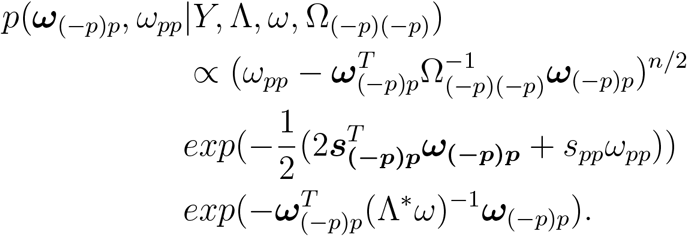

Let

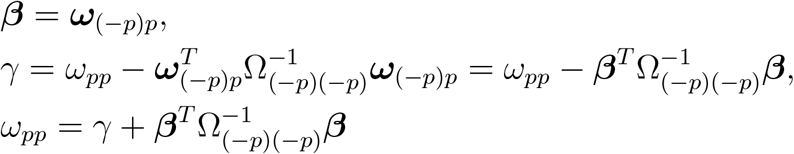

The Jacobian of this transformation is a constant, and the full conditional of *β* and *γ* is

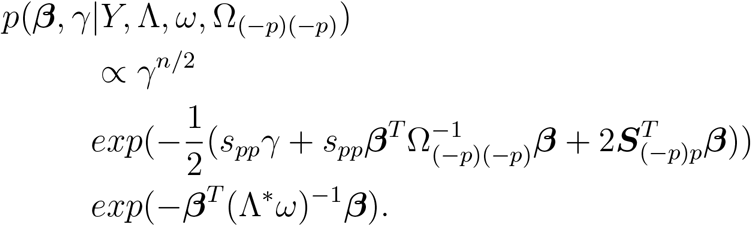

Then, we obtain

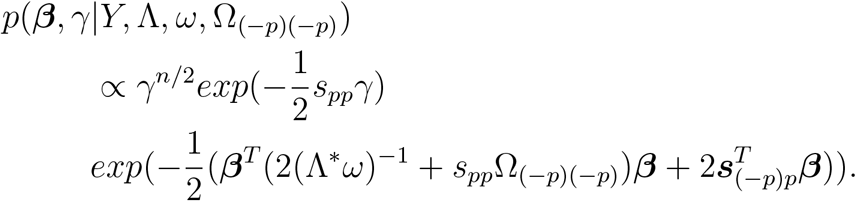

Let *D* = (2(Λ^∗^*ω*)^−1^ + *s*_*pp*_Ω_(−*p*)(−*p*)_)^−1^. Since Ω is positive definite by assumption, we have Ω_(−*p*)(−*p*)_ is symmetric. Hence *D* is symmetric and *D*^*T*^ *D*_−1_ = ***I***. Note that 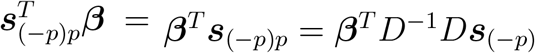 Then, we consider

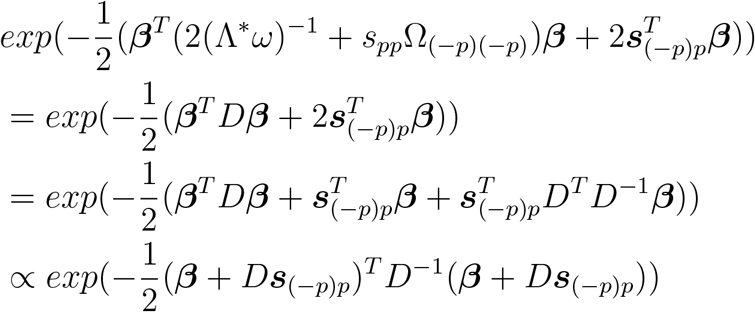

Therefore, we have

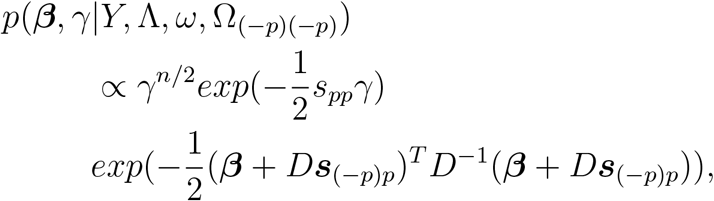

which can be summarized as

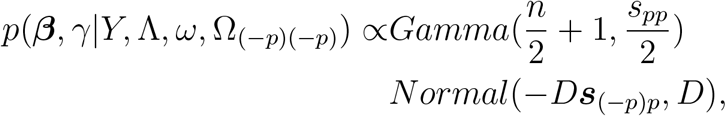

where *D* = (2(Λ^∗^*ω*)^−1^ + *s*_*pp*_Ω_(−*p*)(−*p*)_)^−1^, Λ^∗^ = diag(***λ***_(−*p*)*p*_), and *λ*_*ij*_ = *ϕ*_*ij*_*ψ*_*ij*_.

Above part is about how to sample Ω one column each time. In the following part, we provide standard derivation of the posterior distribution of *ψ*_*ij*_, *ω, ξ* and ***ϕ***.

We first consider 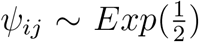 with density function 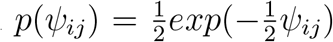. The posterior distribution is

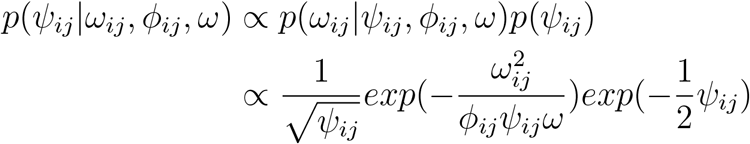

Let 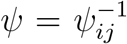, and the Jacobian term is 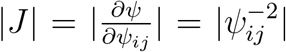. With this transformation, we have

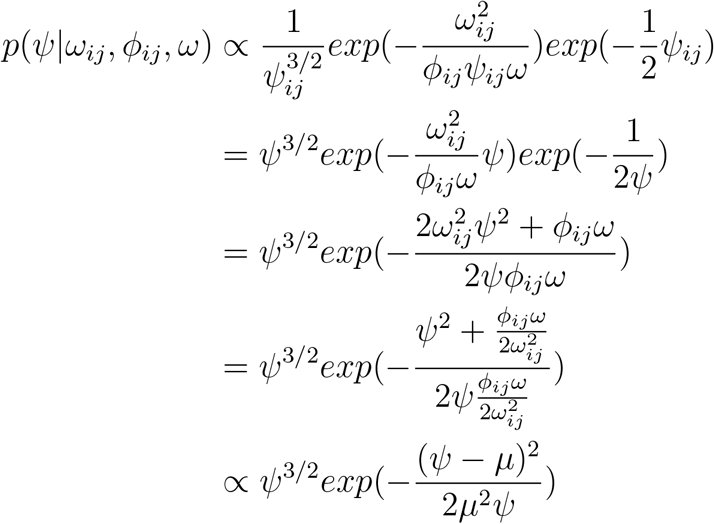

where 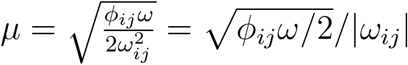 Hence, we can conclude that

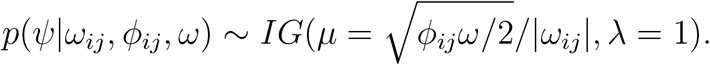

When we sample *ϕ*_*ij*_, we first sample 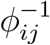 based on above formula and take the reciprocal.

Next we consider *ω*|*ξ* ∼ *Gamma*(*shape* = *a, rate* = *ξ*) with density function 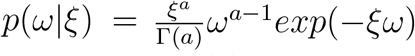 Denote *Z* ∼ giG(*χ, ρ, λ*_0_), the generalized inverse Gaussian distribution, if 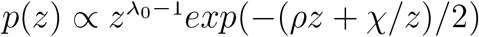. The posterior distribution of *ω* is

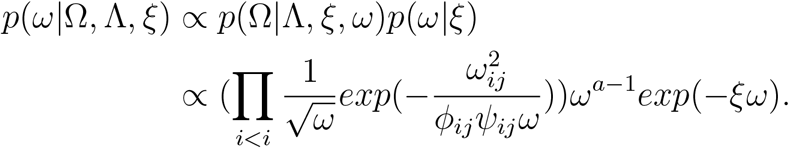

Since we only consider the upper diagonal entries of the precision matrix, there are *p*(*p* − 1)/2 entries in total. We have

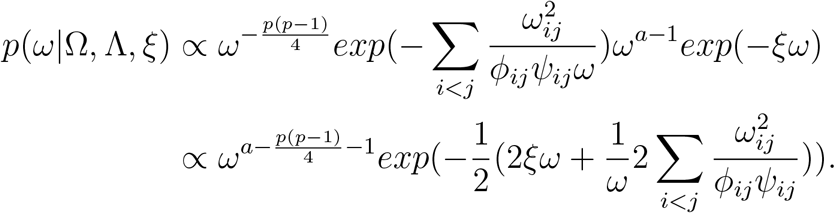

In summary, we have

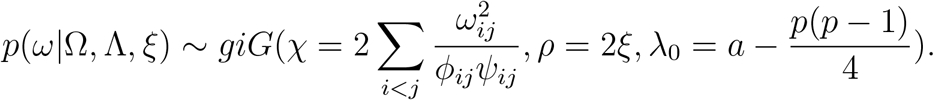

Next, we derive the posterior distribution of *ξ* ∼ *Gamma*(*b*, 1) with probability density function 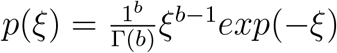.

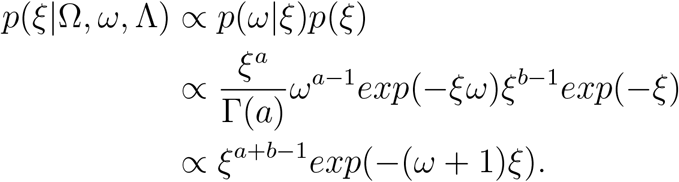

In summary, we have

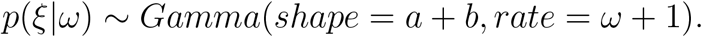

In order to sample ***ϕ*** ∼ *Dir*(*α*_*pi*_, *α*_*pi*_, …, *α*_*pi*_), where there are 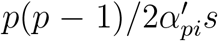 is in total, we resort to the work of [1] and the sampling procedures for ***ϕ*** are in the following. We first integrate out *ω*. The joint distribution of 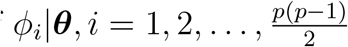 has the form

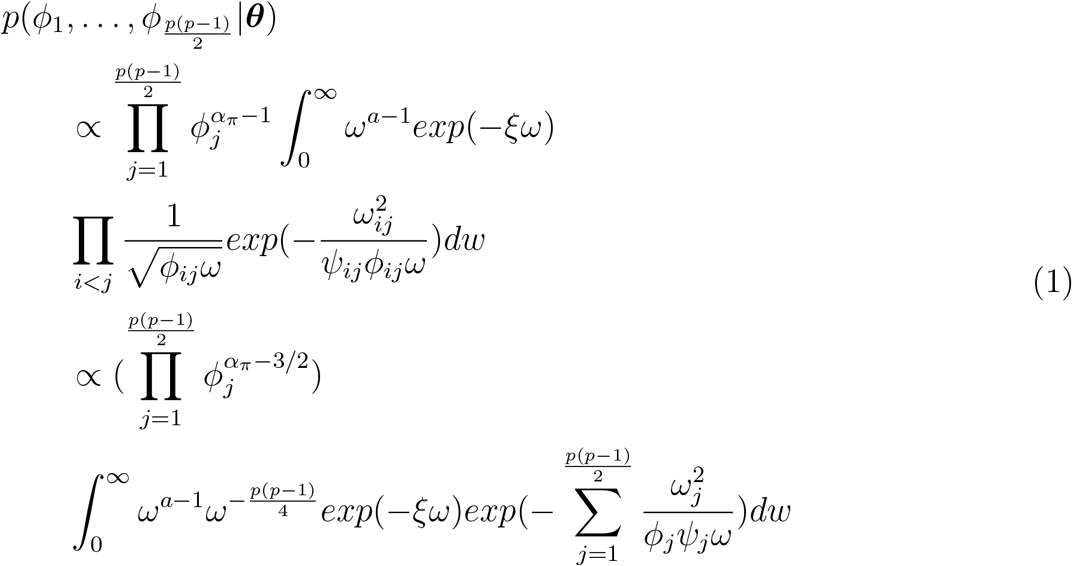

We next use the results from the theory of normalized random measures [2]. Suppose 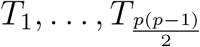 are independent random variables with *T*_*j*_ having a density *f*_*j*_ on (0, +∞). Let *ϕ*_*j*_ = *T*_*j*_/*T* with 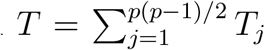. Then, the joint density f of 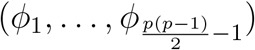 supported on the simplex *S*^*n*−1^ has the form

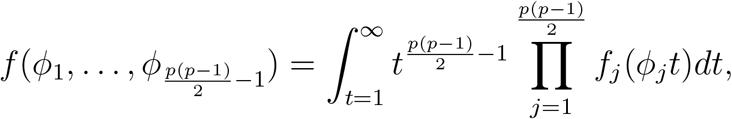

where 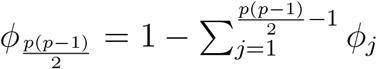. Set

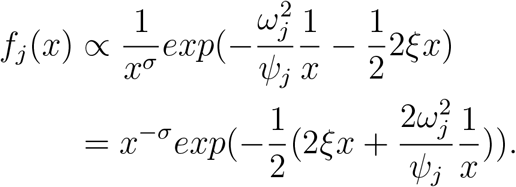

We get

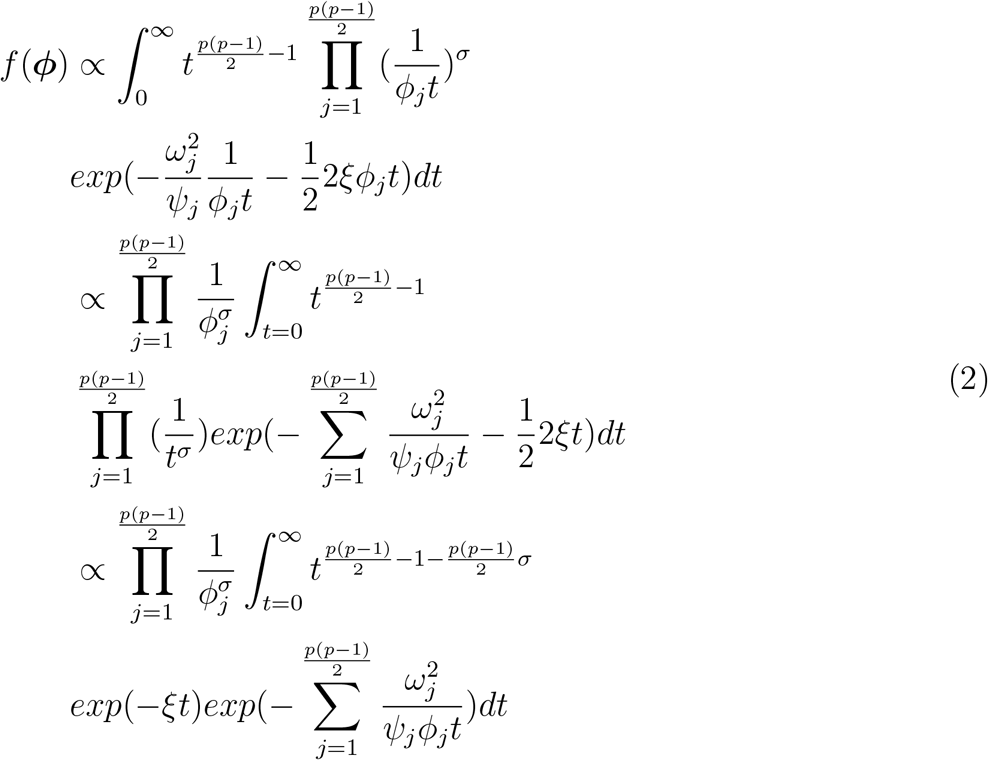

Then, we equate the results from functions 1 and 2 and obtain

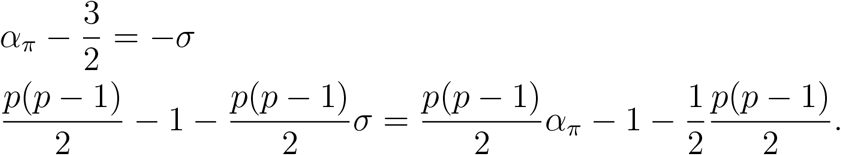

Hence, 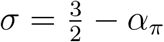 and we have

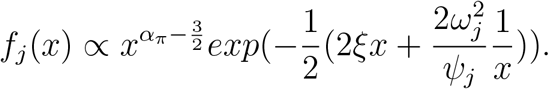

In summary, we have

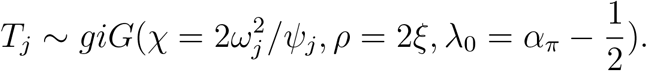

### 2. Formulae of precision, recall, FPR, accuracy and F score

Denote True Positive, False Positive, False Negative and True Negative as TP, FP, FN, and TN respectively. The formulae are

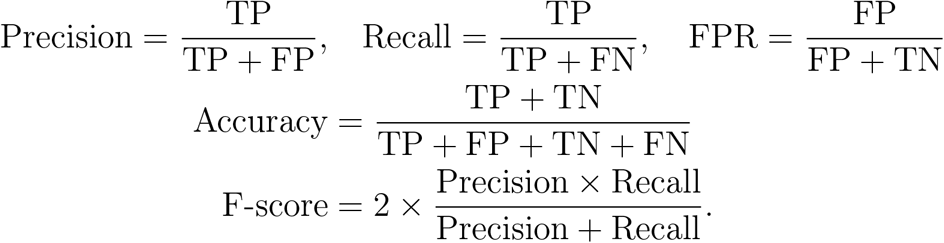

### 3. Supplementary figures

**Figure S1:**
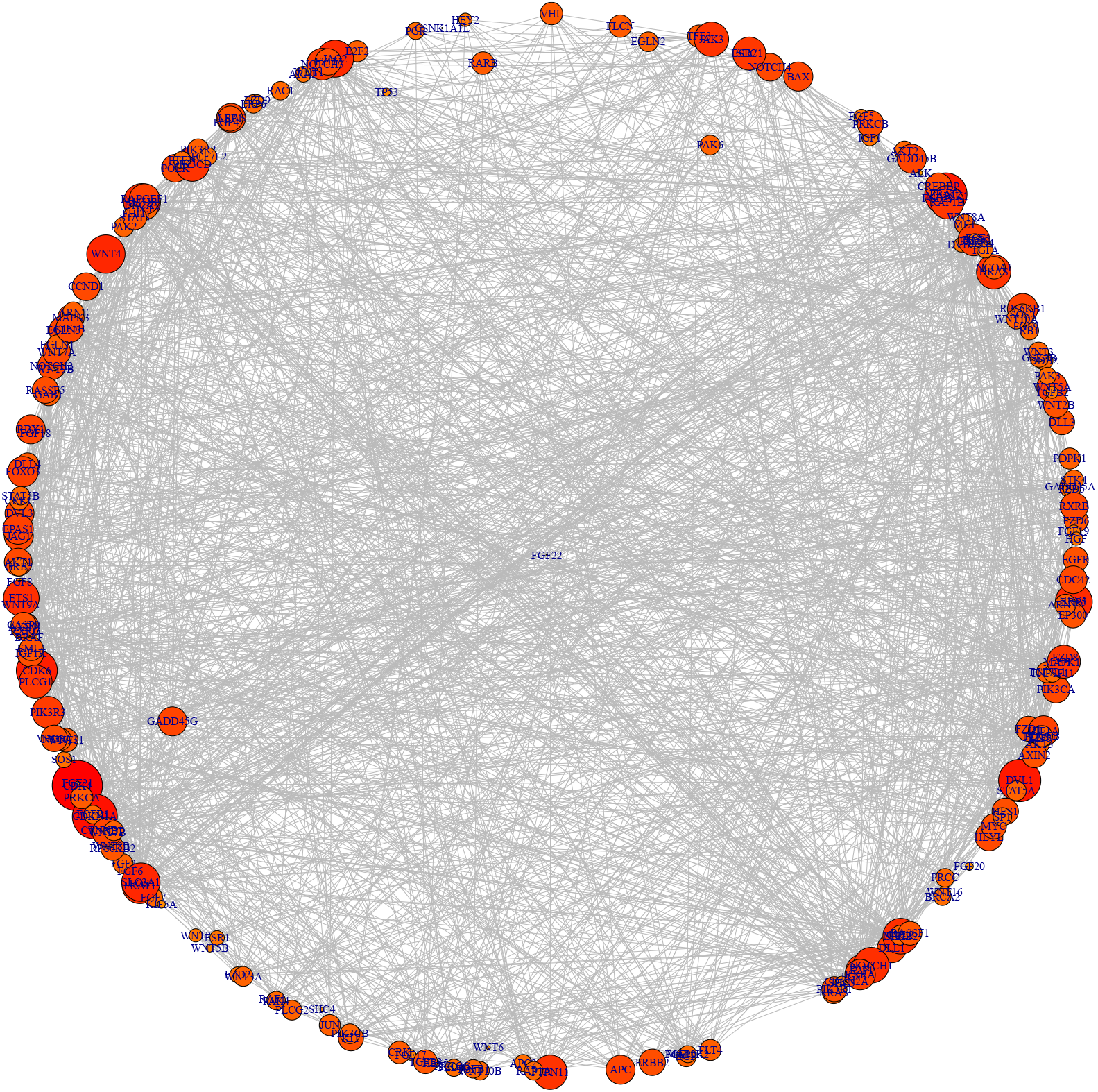
The inferred graph for the LUAD_LUSC dataset by the graphical horseshoe estimator. The size of node is proportional to the degrees of edges.

### 4. Supplementary tables

**Table S1.** Frobenius norm (F-norm), Mean Stein’s loss, precision, recall, false positive rates, accuracy and F scores of precision matrix estimates for network structures Hubs, Cliques positive and Cliques negative over 30 datasets generated by multivariate normal distributions with precision matrix Ω, where p = 100 and n = 50. The precision matrix is estimated by frequentist graphical lasso (GL), the graphical horseshoe (GHS), and the graphical R2D2 (GR2D2). The best performance in each row is shown in bold.

|  | Hubs |  |  | Cliques positive |  |  | Cliques negative |  |  |
| --- | --- | --- | --- | --- | --- | --- | --- | --- | --- |
| (P100, n50) | GR2D2 | GHS | GL | GR2D2 | GHS | GL | GR2D2 | GHS | GL |
| **F-norm** | **3.2188** | 3.4443 | 82.4434 | 5.0073 | **4.869** | 86.3335 | 6.9366 | **6.7698** | 86.7184 |
| (s.d.) | 0.1112 | 0.1385 | 0.5057 | 0.1201 | 0.2189 | 0.667 | 0.0632 | 0.2046 | 0.5267 |
| **Stein's loss** | **5.8591** | 6.7082 | 484.8487 | **53.1149** | 58.6612 | 793.8653 | **23.9461** | 31.6169 | 679.9838 |
| (s.d.) | 0.3417 | 0.4307 | 4.4029 | 2.6451 | 4.7103 | 16.9524 | 1.1757 | 1.9982 | 10.2098 |
| **Precision** | **1** | 0.9981 | 0.0218 | **1** | 0.9969 | 0.0197 | **1** | 0.9989 | 0.0205 |
| (s.d.) | 0 | 0.0059 | 0.0008 | 0 | 0.0074 | 0.0004 | 0 | 0.0044 | 0.0003 |
| **Recall** | 0.5556 | 0.56 | **0.9563** | 0.625 | 0.7967 | **0.9702** | 0.625 | 0.7142 | **0.9962** |
| (s.d.) | 0 | 0.0075 | 0.0267 | 0 | 0.0222 | 0.0157 | 0 | 0.0191 | 0.0068 |
| **FPR** | **0** | 2.04E-05 | 0.7655 | **0** | 4.07E-05 | 0.7650 | **0** | 1.36E-05 | 0.7667 |
| (s.d.) | 0 | 6.21E-05 | 0.0056 | 0 | 9.84E-05 | 0.0053 | 0 | 5.16E-05 | 0.0056 |
| **Accuracy** | 0.992 | **0.9921** | 0.2471 | 0.994 | **0.9967** | 0.2465 | 0.994 | **0.9954** | 0.2454 |
| (s.d.) | 0 | 1.30E-04 | 0.0055 | 0 | 3.59E-04 | 0.0052 | 0 | 3.10E-04 | 0.0055 |
| **F score** | 0.7143 | **7.17E-01** | 0.0427 | 0.7692 | **0.8854** | 0.0387 | 0.7692 | **0.8327** | 0.0402 |
| (s.d.) | 0 | 0.0058 | 0.0015 | 0 | 0.0137 | 0.0007 | 0 | 0.0132 | 0.0005 |

**Table S2.** Frobenius norm (F-norm), Mean Stein’s loss, precision, recall, false positive rates, accuracy and F scores of precision matrix estimates for network structures Hubs, Cliques positive and Cliques negative over 30 datasets generated by multivariate normal distributions with precision matrix Ω, where p = 150 and n = 50. The precision matrix is estimated by frequentist graphical lasso (GL), the graphical horseshoe (GHS), and the graphical R2D2 (GR2D2). The best performance in each row is shown in bold.

|  | Hubs |  |  | Cliques positive |  |  | Cliques negative |  |  |
| --- | --- | --- | --- | --- | --- | --- | --- | --- | --- |
| (P150, n50) | GR2D2 | GHS | GL | GR2D2 | GHS | GL | GR2D2 | GHS | GL |
| **F-norm** | **3.6462** | 4.1748 | 140.6986 | **5.2716** | 5.4002 | 146.4574 | 7.1302 | **7.0766** | 146.1478 |
| (s.d.) | 0.1699 | 0.2507 | 0.3868 | 0.1144 | 0.1894 | 0.6187 | 0.0792 | 0.1833 | 0.628 |
| **Stein's loss** | **6.9167** | 8.7052 | 1154.8461 | **54.249** | 61.8153 | 1631.2601 | **24.4415** | 33.3348 | 1468.3316 |
| (s.d.) | 0.4852 | 0.7139 | 4.7164 | 1.8619 | 3.4329 | 19.6544 | 1.2492 | 1.9639 | 11.2402 |
| **Precision** | **1** | 0.9965 | 0.0150 | **1** | 0.997 | 0.0141 | **1** | 0.9976 | 0.0145 |
| (s.d.) | 0 | 0.0068 | 0.0004 | 0 | 0.0058 | 0.0002 | 0 | 0.0048 | 0.0002 |
| **Recall** | 0.6522 | 0.6545 | **0.9490** | 0.7143 | 0.8435 | **0.9640** | 0.7143 | 0.7911 | **0.9911** |
| (s.d.) | 0 | 0.0051 | 0.0202 | 0 | 0.0192 | 0.0118 | 0 | 0.0214 | 0.009 |
| **FPR** | **0** | 2.39E-05 | 0.6356 | **0** | 2.39E-05 | 0.6315 | **0** | 1.79E-05 | 0.6314 |
| (s.d.) | 0 | 4.68E-05 | 0.0022 | 0 | 4.67E-05 | 0.0029 | 0 | 3.65E-05 | 0.0041 |
| **Accuracy** | **0.9964** | **0.9964** | 0.3703 | 0.9973 | **0.9985** | 0.3740 | 0.9973 | **0.998** | 0.3743 |
| (s.d.) | 0 | 7.38E-05 | 0.0023 | 0 | 1.94E-04 | 0.0029 | 0 | 1.96E-04 | 0.0041 |
| **F score** | 0.7895 | **0.7901** | 0.0295 | 0.8333 | **0.9137** | 0.0277 | 0.8333 | **0.8823** | 0.0285 |
| (s.d.) | 0 | 0.0045 | 0.0008 | 0 | 0.012 | 0.0004 | 0 | 0.0131 | 0.0003 |

**Table S3.** Frobenius norm (F-norm), Mean Stein’s loss, precision, recall, false positive rates, accuracy and F scores of precision matrix estimates for network structures Hubs, Cliques positive and Cliques negative over 30 datasets generated by multivariate normal distributions with precision matrix Ω, where p = 200 and n = 50. The precision matrix is estimated by frequentist graphical lasso (GL), the graphical horseshoe (GHS), and the graphical R2D2 (GR2D2). The best performance in each row is shown in bold.

|  | Hubs |  |  | Cliques positive |  |  | Cliques negative |  |  |
| --- | --- | --- | --- | --- | --- | --- | --- | --- | --- |
| (P200, n50) | GR2D2 | GHS | GL | GR2D2 | GHS | GL | GR2D2 | GHS | GL |
| **F-norm** | **3.9499** | 4.9696 | 191.1832 | **5.5071** | 5.9812 | 197.9015 | **7.3117** | 7.6966 | 197.0806 |
| (s.d.) | 0.1986 | 0.297 | 0.6503 | 0.1106 | 0.2666 | 0.7279 | 0.0932 | 0.311 | 0.6971 |
| **Stein's loss** | **8.0758** | 11.6529 | 1922.5238 | **55.1247** | 64.9922 | 2528.4258 | **26.321** | 38.1823 | 2327.1573 |
| (s.d.) | 0.6396 | 1.055 | 7.8913 | 3.4851 | 4.4964 | 29.9884 | 1.2098 | 2.7868 | 13.0926 |
| **Precision** | **1** | 0.9948 | 0.0123 | **1** | 0.9968 | 0.0117 | **1** | 0.9945 | 0.0120 |
| (s.d.) | 0 | 0.0084 | 0.0003 | 0 | 0.0043 | 0.0002 | 0 | 0.0062 | 0.0002 |
| **Recall** | 0.7143 | 0.7169 | **0.9521** | 0.7692 | 0.8751 | **0.9648** | 0.7692 | 0.8277 | **0.992** |
| (s.d.) | 0 | 0.004 | 0.0156 | 0 | 0.0124 | 0.0106 | 0 | 0.0141 | 0.0066 |
| **FPR** | **0** | 2.69E-05 | 0.5357 | **0** | 1.85E-05 | 0.5311 | **0** | 3.02E-05 | 0.5315 |
| (s.d.) | 0 | 4.33E-05 | 0.0024 | 0 | 2.47E-05 | 0.0026 | 0 | 3.40E-05 | 0.0021 |
| **Accuracy** | **0.998** | **0.998** | 0.4677 | 0.9985 | **0.9992** | 0.4722 | 0.9985 | **0.9989** | 0.4719 |
| (s.d.) | 0 | 5.10E-05 | 0.0024 | 0 | 8.67E-05 | 0.0026 | 0 | 9.19E-05 | 0.0021 |
| **F score** | **0.8333** | **0.8333** | 0.0243 | 0.8696 | **0.932** | 0.0231 | 0.8696 | **0.9034** | 0.0237 |
| (s.d.) | 0 | 0.004 | 0.0005 | 0 | 0.0075 | 0.0003 | 0 | 0.0084 | 0.0003 |

**Table S4.** Frobenius norm (F-norm), Mean Stein’s loss, precision, recall, false positive rates, accuracy and F scores of precision matrix estimates for network structures Hubs, Cliques positive and Cliques negative over 30 datasets generated by multivariate normal distributions with precision matrix Ω, where p = 250 and n = 50. The precision matrix is estimated by frequentist graphical lasso (GL), the graphical horseshoe (GHS), and the graphical R2D2 (GR2D2). The best performance in each row is shown in bold.

|  | Hubs |  |  | Cliques positive |  |  | Cliques negative |  |  |
| --- | --- | --- | --- | --- | --- | --- | --- | --- | --- |
| (P250, n50) | GR2D2 | GHS | GL | GR2D2 | GHS | GL | GR2D2 | GHS | GL |
| **F-norm** | **4.2889** | 5.973 | 236.344 | **5.7492** | 6.816 | 243.6115 | **7.5132** | 8.2696 | 242.1377 |
| (s.d.) | 0.1823 | 0.2517 | 0.657 | 0.1092 | 0.2221 | 0.6882 | 0.1297 | 0.3703 | 0.9657 |
| **Stein's loss** | **9.1185** | 15.6072 | 2745.0562 | **54.7514** | 69.4763 | 3444.198 | **27.779** | 41.837 | 3220.5177 |
| (s.d.) | 0.5655 | 1.038 | 8.2134 | 1.9759 | 3.7656 | 14.67 | 1.9426 | 3.6747 | 15.7963 |
| **Precision** | **1** | 0.9921 | 0.0108 | **1** | 0.9948 | 0.0104 | **1** | 0.9938 | 0.0106 |
| (s.d.) | 0 | 0.0082 | 0.0002 | 0 | 0.0049 | 0.0001 | 0 | 0.0052 | 0.0001 |
| **Recall** | 0.7576 | 0.759 | **0.9518** | 0.8065 | 0.8946 | **0.9700** | 0.8065 | 0.8576 | **0.9889** |
| (s.d.) | 0 | 0.0031 | 0.0168 | 0 | 0.0137 | 0.0097 | 0 | 0.0106 | 0.0084 |
| **FPR** | **0** | 3.22E-05 | 0.4617 | **0** | 2.36E-05 | 0.4575 | **0** | 2.68E-05 | 0.4583 |
| (s.d.) | 0 | 3.38E-05 | 0.0018 | 0 | 2.22E-05 | 0.0013 | 0 | 2.25E-05 | 0.0014 |
| **Accuracy** | **0.9987** | **0.9987** | 0.5405 | 0.999 | **0.9995** | 0.5446 | 0.999 | **0.9993** | 0.5439 |
| (s.d.) | 0 | 3.43E-05 | 0.0018 | 0 | 7.27E-05 | 0.0013 | 0 | 6.02E-05 | 0.0014 |
| **F score** | **0.8621** | 0.86 | 0.0213 | 0.8929 | **0.942** | 0.0207 | 0.8929 | **0.9207** | 0.0211 |
| (s.d.) | 0 | 0.0033 | 0.0004 | 0 | 0.0081 | 0.0002 | 0 | 0.0069 | 0.0002 |

**Table S5.** Frobenius norm (F-norm), Mean Stein’s loss, precision, recall, false positive rates, accuracy and F scores of precision matrix estimates for scale-free network structures Positive, Negative and Half over 30 datasets generated by multivariate normal distributions with precision matrix Ω, where p = 100 and n = 50. The precision matrix is estimated by frequentist graphical lasso (GL), the graphical horseshoe (GHS), and the graphical R2D2 (GR2D2). The best performance in each row is shown in bold.

|  | Scale-free positive |  |  | Scale-free negative |  |  | Scale-free half |  |  |
| --- | --- | --- | --- | --- | --- | --- | --- | --- | --- |
| (P100, n50) | GR2D2 | GHS | GL | GR2D2 | GHS | GL | GR2D2 | GHS | GL |
| **F-norm** | **3.0680** | 3.2880 | 82.4385 | **3.0928** | 3.3613 | 82.5090 | **3.1336** | 3.3995 | 82.6982 |
| (s.d.) | 0.1308 | 0.2082 | 0.5793 | 0.1814 | 0.2342 | 0.5100 | 0.1245 | 0.1798 | 0.4436 |
| **Stein's loss** | **4.8533** | 5.5368 | 480.9690 | **4.9342** | 5.7451 | 482.3797 | **5.0948** | 5.8694 | 483.5125 |
| (s.d.) | 0.3414 | 0.5583 | 4.8474 | 0.4550 | 0.5823 | 4.3023 | 0.3007 | 0.4279 | 3.7743 |
| **Precision** | **1** | 0.9981 | 0.0312 | **1** | 0.9961 | 0.0313 | **1** | 0.9967 | 0.0310 |
| (s.d.) | 0 | 0.0078 | 0.0012 | 0 | 0.0080 | 0.0012 | 0 | 0.0074 | 0.0010 |
| **Recall** | 0.3356 | 0.3358 | **0.8826** | 0.3356 | 0.3362 | **0.8864** | 0.3356 | 0.3360 | **0.8856** |
| (s.d.) | 0 | 0.0012 | 0.0258 | 0 | 0.0020 | 0.0244 | 0 | 0.0017 | 0.0230 |
| **FPR** | **0** | 2.0614E-05 | 0.7646 | **0** | 4.1229E-05 | 0.7641 | **0** | 3.4357E-05 | 0.7629 |
| (s.d.) | 0 | 8.2989E-05 | 0.0056 | 0 | 8.3867E-05 | 0.0051 | 0 | 7.8138E-05 | 0.0047 |
| **Accuracy** | **0.9802** | **0.9802** | 0.2530 | **0.9802** | **0.9802** | 0.2535 | **0.9802** | **0.9802** | 0.2544 |
| (s.d.) | 0 | 8.9955E-05 | 0.0055 | 0 | 9.6132E-05 | 0.0052 | 0 | 9.6132E-05 | 0.0046 |
| **F score** | **0.5025** | **0.5025** | 0.0602 | 0.5025 | **0.5028** | 0.0605 | 0.5025 | **0.5026** | 0.0599 |
| (s.d.) | 0 | 0.0017 | 0.0024 | 0 | 0.0024 | 0.0022 | 0 | 0.0022 | 0.0018 |

**Table S6.**
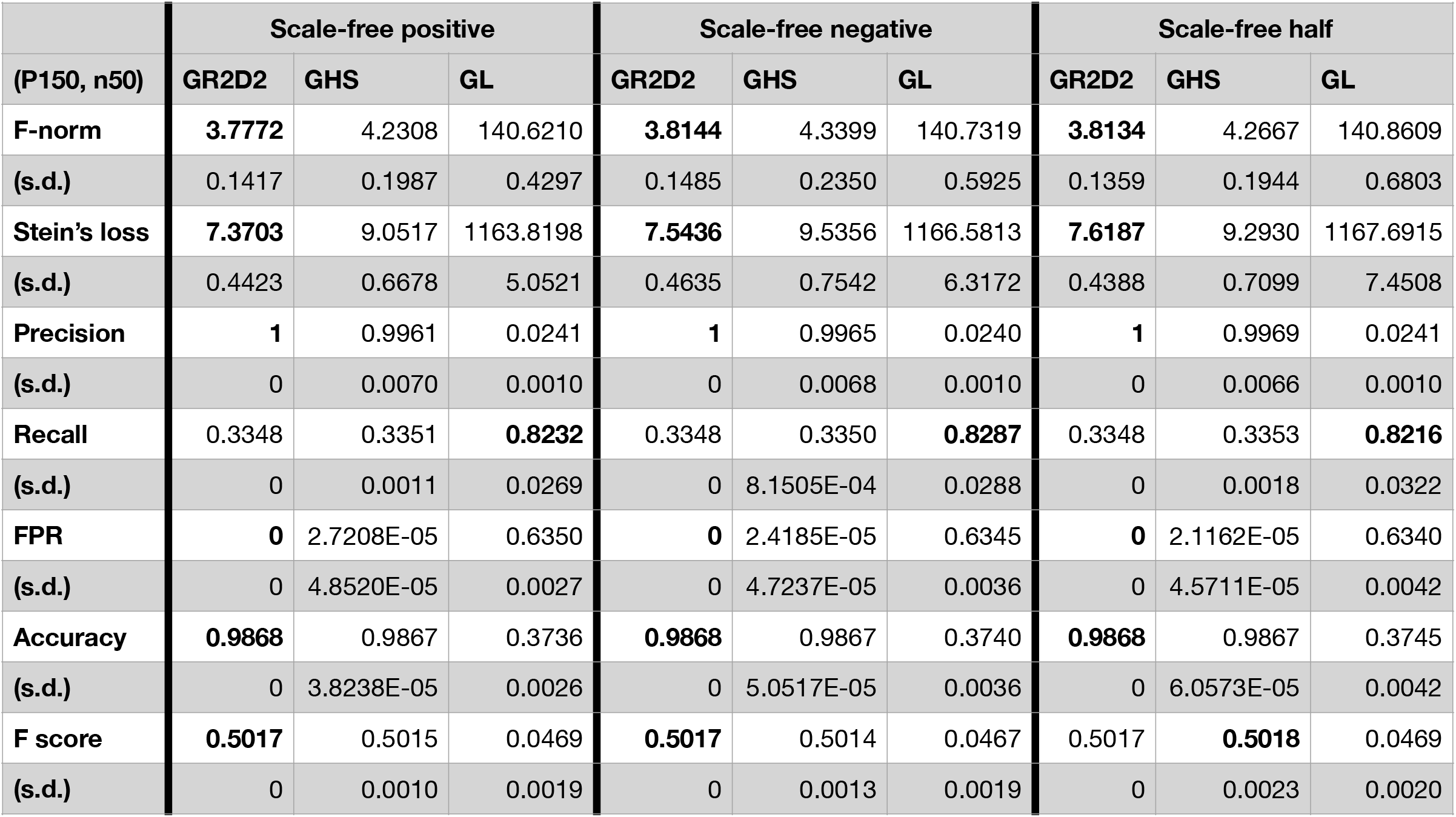
Frobenius norm (F-norm), Mean Stein’s loss, precision, recall, false positive rates, accuracy and F scores of precision matrix estimates for scale-free network structures Positive, Negative and Half over 30 datasets generated by multivariate normal distributions with precision matrix Ω, where p = 150 and n = 50. The precision matrix is estimated by frequentist graphical lasso (GL), the graphical horseshoe (GHS), and the graphical R2D2 (GR2D2). The best performance in each row is shown in bold.

|  | Scale-free positive |  |  | Scale-free negative |  |  | Scale-free half |  |  |
| --- | --- | --- | --- | --- | --- | --- | --- | --- | --- |
| (P150, n50) | GR2D2 | GHS | GL | GR2D2 | GHS | GL | GR2D2 | GHS | GL |
| **F-norm** | **3.7772** | 4.2308 | 140.6210 | **3.8144** | 4.3399 | 140.7319 | **3.8134** | 4.2667 | 140.8609 |
| (s.d.) | 0.1417 | 0.1987 | 0.4297 | 0.1485 | 0.2350 | 0.5925 | 0.1359 | 0.1944 | 0.6803 |
| **Stein's loss** | **7.3703** | 9.0517 | 1163.8198 | **7.5436** | 9.5356 | 1166.5813 | **7.6187** | 9.2930 | 1167.6915 |
| (s.d.) | 0.4423 | 0.6678 | 5.0521 | 0.4635 | 0.7542 | 6.3172 | 0.4388 | 0.7099 | 7.4508 |
| **Precision** | **1** | 0.9961 | 0.0241 | **1** | 0.9965 | 0.0240 | **1** | 0.9969 | 0.0241 |
| (s.d.) | 0 | 0.0070 | 0.0010 | 0 | 0.0068 | 0.0010 | 0 | 0.0066 | 0.0010 |
| **Recall** | 0.3348 | 0.3351 | **0.8232** | 0.3348 | 0.3350 | **0.8287** | 0.3348 | 0.3353 | **0.8216** |
| (s.d.) | 0 | 0.0011 | 0.0269 | 0 | 8.1505E-04 | 0.0288 | 0 | 0.0018 | 0.0322 |
| **FPR** | **0** | 2.7208E-05 | 0.6350 | **0** | 2.4185E-05 | 0.6345 | **0** | 2.1162E-05 | 0.6340 |
| (s.d.) | 0 | 4.8520E-05 | 0.0027 | 0 | 4.7237E-05 | 0.0036 | 0 | 4.5711E-05 | 0.0042 |
| **Accuracy** | **0.9868** | 0.9867 | 0.3736 | **0.9868** | 0.9867 | 0.3740 | **0.9868** | 0.9867 | 0.3745 |
| (s.d.) | 0 | 3.8238E-05 | 0.0026 | 0 | 5.0517E-05 | 0.0036 | 0 | 6.0573E-05 | 0.0042 |
| **F score** | **0.5017** | 0.5015 | 0.0469 | **0.5017** | 0.5014 | 0.0467 | 0.5017 | **0.5018** | 0.0469 |
| (s.d.) | 0 | 0.0010 | 0.0019 | 0 | 0.0013 | 0.0019 | 0 | 0.0023 | 0.0020 |

**Table S7.**
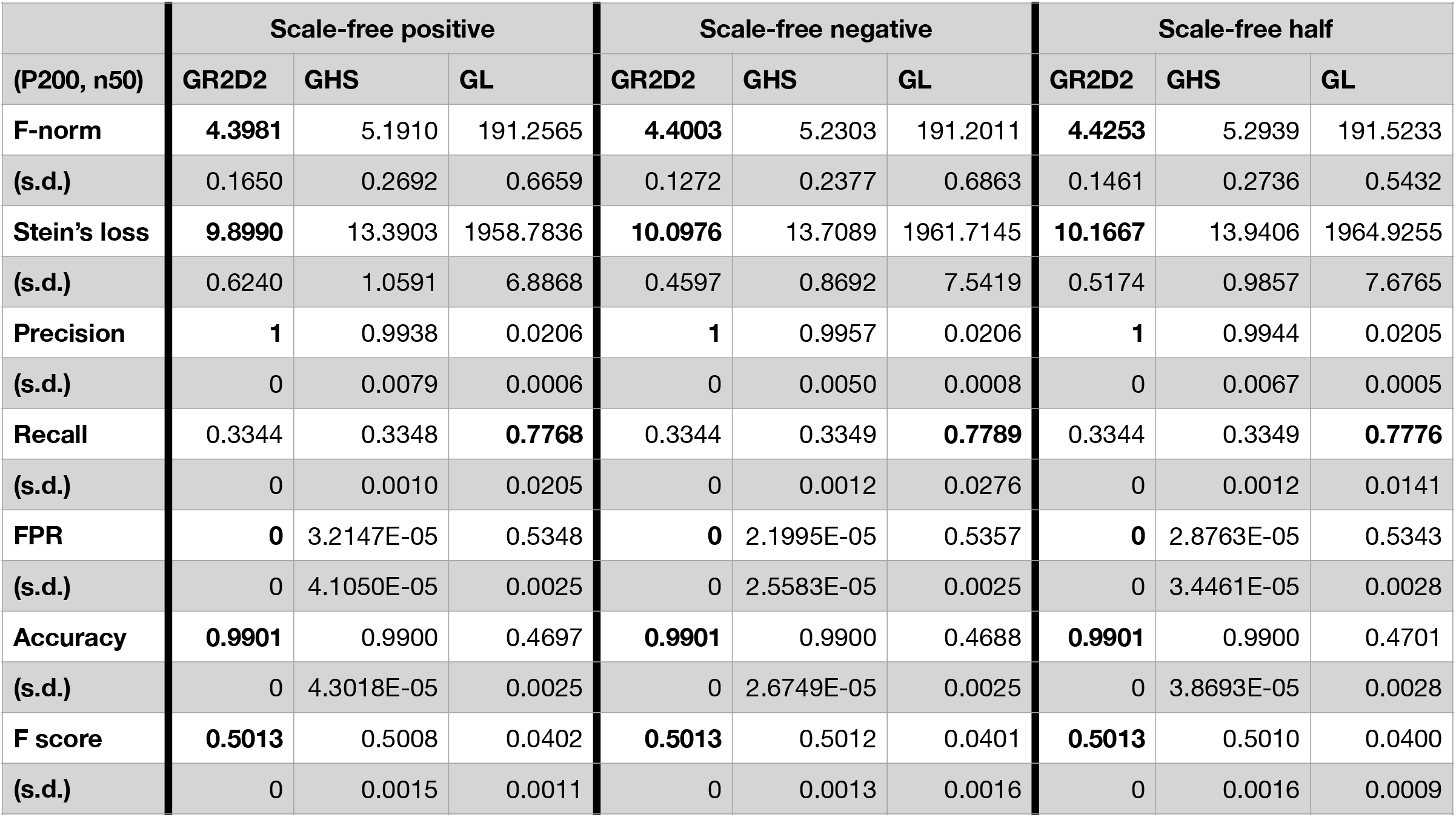
Frobenius norm (F-norm), Mean Stein’s loss, precision, recall, false positive rates, accuracy and F scores of precision matrix estimates for scale-free network structures Positive, Negative and Half over 30 datasets generated by multivariate normal distributions with precision matrix Ω, where p = 200 and n = 50. The precision matrix is estimated by frequentist graphical lasso (GL), the graphical horseshoe (GHS), and the graphical R2D2 (GR2D2). The best performance in each row is shown in bold.

|  | Scale-free positive |  |  | Scale-free negative |  |  | Scale-free half |  |  |
| --- | --- | --- | --- | --- | --- | --- | --- | --- | --- |
| (P200, n50) | GR2D2 | GHS | GL | GR2D2 | GHS | GL | GR2D2 | GHS | GL |
| **F-norm** | **4.3981** | 5.1910 | 191.2565 | **4.4003** | 5.2303 | 191.2011 | **4.4253** | 5.2939 | 191.5233 |
| (s.d.) | 0.1650 | 0.2692 | 0.6659 | 0.1272 | 0.2377 | 0.6863 | 0.1461 | 0.2736 | 0.5432 |
| **Stein's loss** | **9.8990** | 13.3903 | 1958.7836 | **10.0976** | 13.7089 | 1961.7145 | **10.1667** | 13.9406 | 1964.9255 |
| (s.d.) | 0.6240 | 1.0591 | 6.8868 | 0.4597 | 0.8692 | 7.5419 | 0.5174 | 0.9857 | 7.6765 |
| **Precision** | **1** | 0.9938 | 0.0206 | **1** | 0.9957 | 0.0206 | **1** | 0.9944 | 0.0205 |
| (s.d.) | 0 | 0.0079 | 0.0006 | 0 | 0.0050 | 0.0008 | 0 | 0.0067 | 0.0005 |
| **Recall** | 0.3344 | 0.3348 | **0.7768** | 0.3344 | 0.3349 | **0.7789** | 0.3344 | 0.3349 | **0.7776** |
| (s.d.) | 0 | 0.0010 | 0.0205 | 0 | 0.0012 | 0.0276 | 0 | 0.0012 | 0.0141 |
| **FPR** | **0** | 3.2147E-05 | 0.5348 | **0** | 2.1995E-05 | 0.5357 | **0** | 2.8763E-05 | 0.5343 |
| (s.d.) | 0 | 4.1050E-05 | 0.0025 | 0 | 2.5583E-05 | 0.0025 | 0 | 3.4461E-05 | 0.0028 |
| **Accuracy** | **0.9901** | 0.9900 | 0.4697 | **0.9901** | 0.9900 | 0.4688 | **0.9901** | 0.9900 | 0.4701 |
| (s.d.) | 0 | 4.3018E-05 | 0.0025 | 0 | 2.6749E-05 | 0.0025 | 0 | 3.8693E-05 | 0.0028 |
| **F score** | **0.5013** | 0.5008 | 0.0402 | **0.5013** | 0.5012 | 0.0401 | **0.5013** | 0.5010 | 0.0400 |
| (s.d.) | 0 | 0.0015 | 0.0011 | 0 | 0.0013 | 0.0016 | 0 | 0.0016 | 0.0009 |

**Table S8.** Frobenius norm (F-norm), Mean Stein’s loss, precision, recall, false positive rates, accuracy and F scores of precision matrix estimates for scale-free network structures Positive, Negative and Half over 30 datasets generated by multivariate normal distributions with precision matrix Ω, where p = 250 and n = 50. The precision matrix is estimated by frequentist graphical lasso (GL), the graphical horseshoe (GHS), and the graphical R2D2 (GR2D2). The best performance in each row is shown in bold.

|  | Scale-free positive |  |  | Scale-free negative |  |  | Scale-free half |  |  |
| --- | --- | --- | --- | --- | --- | --- | --- | --- | --- |
| (P250, n50) | GR2D2 | GHS | GL | GR2D2 | GHS | GL | GR2D2 | GHS | GL |
| **F-norm** | **5.0374** | 6.2186 | 236.6587 | **5.0073** | 6.2395 | 236.1609 | **5.0266** | 6.3137 | 236.6091 |
| (s.d.) | 0.1091 | 0.3048 | 0.6145 | 0.0174 | 0.2449 | 0.8115 | 0.2099 | 0.3806 | 0.6069 |
| **Stein's loss** | **12.9199** | 18.8915 | 2819.6789 | **12.9766** | 19.2947 | 2817.7321 | **13.0276** | 19.4635 | 2822.2823 |
| (s.d.) | 0.3989 | 1.2837 | 6.0606 | 0.4700 | 1.2020 | 11.9869 | 0.8417 | 1.7436 | 7.2261 |
| **Precision** | **1** | 0.9927 | 0.0187 | **1** | 0.9937 | 0.0187 | **1** | 0.9911 | 0.0185 |
| (s.d.) | 0 | 0.0086 | 0.0006 | 0 | 0.0063 | 0.0004 | 0 | 0.0070 | 0.0003 |
| **Recall** | 0.3342 | 0.3348 | **0.7505** | 0.3342 | 0.3350 | **0.7564** | 0.3342 | 0.3346 | **0.7487** |
| (s.d.) | 0 | 0.0012 | 0.0217 | 0 | 0.0016 | 0.0169 | 0 | 9.2445E-04 | 0.0157 |
| **FPR** | **0** | 3.0228E-05 | 0.4597 | **0** | 2.5910E-05 | 0.4621 | **0** | 3.6706E-05 | 0.4615 |
| (s.d.) | 0 | 3.6018E-05 | 0.0023 | 0 | 2.6077E-05 | 0.0018 | 0 | 2.9134E-05 | 0.0022 |
| **Accuracy** | **0.9920** | 0.9920 | 0.5427 | **0.9920** | **0.9920** | 0.5404 | **0.9920** | **0.9920** | 0.5410 |
| (s.d.) | 0 | 2.8061E-05 | 0.0024 | 0 | 3.1161E-05 | 0.0017 | 0 | 2.9111E-05 | 0.0021 |
| **F score** | **0.5010** | 0.5008 | 0.0364 | 0.5010 | **0.5011** | 0.0365 | **0.5010** | 0.5003 | 0.0361 |
| (s.d.) | 0 | 9.7054E-04 | 0.0011 | 0 | 0.0019 | 0.0008 | 0 | 0.0013 | 0.0006 |

**Table S9.**
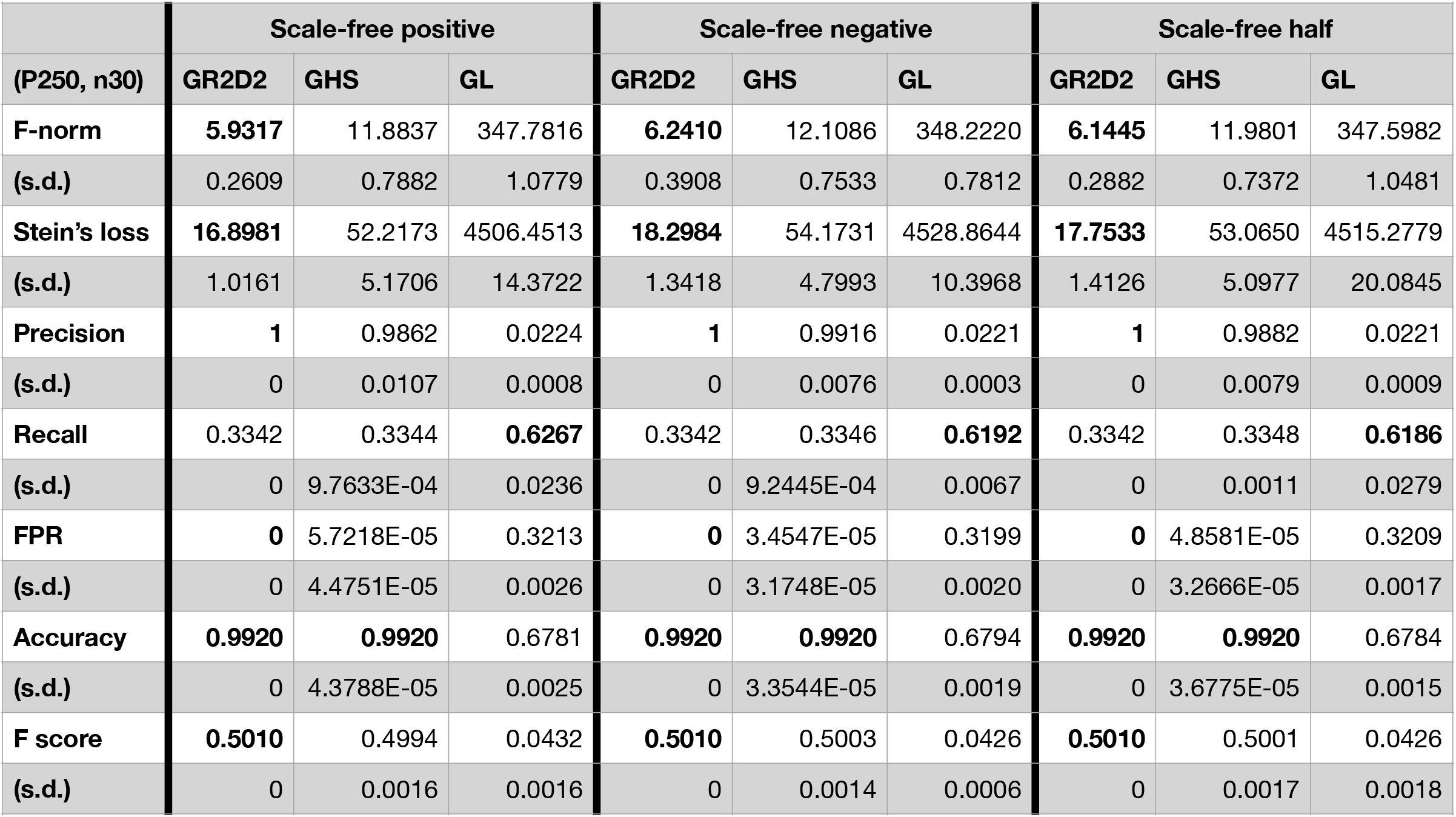
Frobenius norm (F-norm), Mean Stein’s loss, precision, recall, false positive rates, accuracy and F scores of precision matrix estimates for scale-free network structures Positive, Negative and Half over 30 datasets generated by multivariate normal distributions with precision matrix Ω, where p = 250 and n = 30. The precision matrix is estimated by frequentist graphical lasso (GL), the graphical horseshoe (GHS), and the graphical R2D2 (GR2D2). The best performance in each row is shown in bold.

|  | Scale-free positive |  |  | Scale-free negative |  |  | Scale-free half |  |  |
| --- | --- | --- | --- | --- | --- | --- | --- | --- | --- |
| (P250, n30) | GR2D2 | GHS | GL | GR2D2 | GHS | GL | GR2D2 | GHS | GL |
| **F-norm** | **5.9317** | 11.8837 | 347.7816 | **6.2410** | 12.1086 | 348.2220 | **6.1445** | 11.9801 | 347.5982 |
| (s.d.) | 0.2609 | 0.7882 | 1.0779 | 0.3908 | 0.7533 | 0.7812 | 0.2882 | 0.7372 | 1.0481 |
| **Stein's loss** | **16.8981** | 52.2173 | 4506.4513 | **18.2984** | 54.1731 | 4528.8644 | **17.7533** | 53.0650 | 4515.2779 |
| (s.d.) | 1.0161 | 5.1706 | 14.3722 | 1.3418 | 4.7993 | 10.3968 | 1.4126 | 5.0977 | 20.0845 |
| **Precision** | **1** | 0.9862 | 0.0224 | **1** | 0.9916 | 0.0221 | **1** | 0.9882 | 0.0221 |
| (s.d.) | 0 | 0.0107 | 0.0008 | 0 | 0.0076 | 0.0003 | 0 | 0.0079 | 0.0009 |
| **Recall** | 0.3342 | 0.3344 | **0.6267** | 0.3342 | 0.3346 | **0.6192** | 0.3342 | 0.3348 | **0.6186** |
| (s.d.) | 0 | 9.7633E-04 | 0.0236 | 0 | 9.2445E-04 | 0.0067 | 0 | 0.0011 | 0.0279 |
| **FPR** | **0** | 5.7218E-05 | 0.3213 | **0** | 3.4547E-05 | 0.3199 | **0** | 4.8581E-05 | 0.3209 |
| (s.d.) | 0 | 4.4751E-05 | 0.0026 | 0 | 3.1748E-05 | 0.0020 | 0 | 3.2666E-05 | 0.0017 |
| **Accuracy** | **0.9920** | **0.9920** | 0.6781 | **0.9920** | **0.9920** | 0.6794 | **0.9920** | **0.9920** | 0.6784 |
| (s.d.) | 0 | 4.3788E-05 | 0.0025 | 0 | 3.3544E-05 | 0.0019 | 0 | 3.6775E-05 | 0.0015 |
| **F score** | **0.5010** | 0.4994 | 0.0432 | **0.5010** | 0.5003 | 0.0426 | **0.5010** | 0.5001 | 0.0426 |
| (s.d.) | 0 | 0.0016 | 0.0016 | 0 | 0.0014 | 0.0006 | 0 | 0.0017 | 0.0018 |

**Table S10.** Frobenius norm (F-norm), Mean Stein’s loss, precision, recall, false positive rates, accuracy and F scores of precision matrix estimates for network structures Hubs, Cliques positive and Cliques negative over 30 different datasets generated by multivariate normal distributions with 30 different precision matrices Ω, where p = 100 and n = 50. The precision matrix is estimated by frequentist graphical lasso (GL), the graphical horseshoe (GHS), and the graphical R2D2 (GR2D2). The best performance in each row is shown in bold.

|  | Different.Hubs |  |  | Different.Cliques positive |  |  | Different.Cliques negative |  |  |
| --- | --- | --- | --- | --- | --- | --- | --- | --- | --- |
| (P100, n50) | GR2D2 | GHS | GL | GR2D2 | GHS | GL | GR2D2 | GHS | GL |
| **F-norm** | **3.4594** | 3.6141 | 82.9108 | **3.1724** | 3.4886 | 82.8688 | **3.0308** | 3.3812 | 82.5949 |
| **(s.d.)** | 0.8119 | 0.6405 | **0.6288** | 0.6422 | **0.4880** | 0.7030 | 0.4422 | 0.4894 | **0.3415** |
| **Stein's loss** | **8.6314** | 9.3594 | 501.4684 | **7.6612** | 8.8059 | 494.3325 | **4.0098** | 4.9211 | 478.5753 |
| **(s.d.)** | **6.0021** | 6.3452 | 33.4307 | 5.3531 | **5.2763** | 28.2741 | **1.0906** | 1.3944 | 8.7506 |
| **Precision** | **1** | 0.9934 | 0.02201 | **1** | 0.9969 | 0.0196 | **1** | 0.9968 | 0.0191 |
| **(s.d.)** | **0** | 0.0152 | 0.0010 | **0** | 0.0070 | 0.0007 | **0** | 0.0089 | 0.0013 |
| **Recall** | 0.5556 | 0.5726 | **0.9585** | 0.625 | 0.6550 | **0.9597** | 0.625 | 0.6267 | **0.9476** |
| **(s.d.)** | **0** | 0.0265 | 0.0263 | **0** | 0.0397 | 0.0196 | **0** | 0.0043 | 0.0355 |
| **FPR** | **0** | 7.47E-05 | 0.7628 | **0** | 3.39E-05 | 0.7649 | **0** | 3.39E-05 | 0.7653 |
| **(s.d.)** | **0** | 1.73E-04 | 0.0057 | **0** | 7.70E-05 | 0.0037 | **0** | 9.37E-05 | 0.0054 |
| **Accuracy** | 0.992 | **0.9922** | 0.2498 | 0.994 | **0.9944** | 0.2465 | **0.994** | **0.9940** | 0.2457 |
| **(s.d.)** | **0** | 4.58E-04 | 0.0056 | **0** | 6.25E-04 | 0.0038 | **0** | 1.23E-04 | 0.0056 |
| **F score** | 0.7143 | **7.26E-01** | 0.0430 | 0.7692 | **0.7899** | 0.0384 | 0.7692 | **0.7695** | 0.0373 |
| **(s.d.)** | **0** | 0.0200 | 0.0020 | **0** | 0.0277 | 0.0014 | **0** | 0.0045 | 0.0025 |

**Table S11.** Frobenius norm (F-norm), Mean Stein’s loss, precision, recall, false positive rates, accuracy and F scores of precision matrix estimates for network structures Hubs, Cliques positive and Cliques negative over 30 different datasets generated by multivariate normal distributions with 30 different precision matrices Ω, where p = 150 and n = 50. The precision matrix is estimated by frequentist graphical lasso (GL), the graphical horseshoe (GHS), and the graphical R2D2 (GR2D2). The best performance in each row is shown in bold.

|  | Different.Hubs |  |  | Different.Cliques positive |  |  | Different.Cliques negative |  |  |
| --- | --- | --- | --- | --- | --- | --- | --- | --- | --- |
| (P150, n50) | GR2D2 | GHS | GL | GR2D2 | GHS | GL | GR2D2 | GHS | GL |
| **F-norm** | **3.8987** | 4.3683 | 141.2842 | **3.7038** | 4.2616 | 141.2350 | **3.6989** | 4.1376 | 140.8548 |
| **(s.d.)** | 0.7092 | 0.6009 | **0.5567** | 0.6311 | 0.4984 | **0.4553** | 0.4910 | 0.4638 | **0.4384** |
| **Stein's loss** | **10.1162** | 11.5502 | 1184.3517 | **9.7600** | 11.4109 | 1175.2010 | **5.8451** | 7.1299 | 1148.7890 |
| **(s.d.)** | 6.5300 | **6.4673** | 58.1842 | **5.9547** | 6.2515 | 53.1417 | **1.4292** | 1.6567 | 17.7214 |
| **Precision** | **1** | 0.9904 | 0.0151 | **1** | 0.9949 | 0.0138 | **1** | 0.9969 | 0.0136 |
| **(s.d.)** | **0** | 0.0103 | 0.0006 | **0** | 0.0071 | 0.0006 | **0** | 0.0056 | 0.0008 |
| **Recall** | 0.6522 | 0.6672 | **0.9528** | 0.7143 | 0.7368 | **0.9554** | 0.7143 | 0.7149 | **0.9482** |
| **(s.d.)** | **0** | 0.0245 | 0.0278 | **0** | 0.0314 | 0.0279 | **0** | 0.0024 | 0.0400 |
| **FPR** | **0** | 6.89E-05 | 0.6342 | **0** | 3.59E-05 | 0.6339 | **0** | 2.09E-05 | 0.6339 |
| **(s.d.)** | **0** | 7.71E-05 | 0.0033 | **0** | 5.05E-05 | 0.0038 | **0** | 3.86E-05 | 0.0021 |
| **Accuracy** | 0.9964 | **0.9965** | 0.3717 | 0.9973 | **0.9975** | 0.3715 | **0.9973** | **0.9973** | 0.3714 |
| **(s.d.)** | **0** | 2.08E-04 | 0.0033 | **0** | 2.95E-04 | 0.0036 | **0** | 4.10E-05 | 0.0023 |
| **F score** | 0.7895 | **0.7970** | 0.0297 | 0.8333 | **0.8463** | 0.0273 | **0.8333** | 0.8327 | 0.0269 |
| **(s.d.)** | **0** | 0.0152 | 0.0011 | **0** | 0.0204 | 0.0012 | **0** | 0.0023 | 0.0016 |

